# A memory of RPS25 loss drives resistance phenotypes

**DOI:** 10.1101/805663

**Authors:** Alex G. Johnson, Ryan A. Flynn, Christopher P. Lapointe, Yaw Shin Ooi, Michael L. Zhao, Christopher M. Richards, Wenjie Qiao, Shizuka B. Yamada, Julien Couthouis, Aaron D. Gitler, Jan E. Carette, Joseph D. Puglisi

## Abstract

In order to maintain cellular protein homeostasis, ribosomes are safeguarded against dysregulation by myriad processes. Many cell types can nonetheless withstand genetic lesions of certain ribosomal protein genes, some of which are linked to diverse cellular phenotypes and human disease. However, the direct and indirect consequences from sustained alterations in ribosomal protein levels are poorly understood. To address this knowledge gap, we studied *in vitro* and cellular consequences that follow genetic knockout of the ribosomal proteins RPS25 or RACK1 in a human cell line, as both proteins are implicated in direct translational control. Prompted by the unexpected detection of an off-target ribosome alteration in the RPS25 knockout, we closely interrogated cellular phenotypes. We found that multiple RPS25 knockout clones display viral- and toxin-resistance phenotypes that cannot be rescued by functional cDNA expression, suggesting that RPS25 loss elicits a cell state transition. We characterized this state and found that it underlies pleiotropic phenotypes and has a common rewiring of gene expression. Rescuing RPS25 expression by genomic locus repair failed to correct for the phenotypic and expression hysteresis. Our findings illustrate how the elasticity of cells to a ribosome perturbation can drive specific phenotypic outcomes that are indirectly linked to translation.

## INTRODUCTION

The eukaryotic ribosome is comprised of four strands of rRNA and ∼80 ribosomal proteins (RPs), most of which are essential for life. To ensure accurate and efficient protein synthesis, cells have evolved numerous measures to control and protect the cellular ribosome pool. The existence of genetic knockouts of select RPs in yeast and human cell lines nevertheless indicates that cells are elastic to ribosome compositional alterations (1, 2). The presence of ribosomes with substoichiometric RP levels in unperturbed cells further suggests the possibility that certain alterations might represent direct, regulated control of protein synthesis by RPs (3, 4). However, alterations could also represent ribosomes that have escaped from imperfect cellular quality control measures. While not eliciting cell death, RP alteration might be sensed and lead to diverse indirect cellular outcomes. RP loss may therefore drive both direct effects on translation and indirect effects as cells sense and adapt to ribosome irregularities. While genetic RP loss is linked to numerous cellular phenotypes and human disease, the mechanistic basis by which these alterations arise remains unclear (5).

For RP composition of the ribosome to control protein synthesis directly, specific molecular interactions between ribosome-bound RPs and mRNA transcripts may occur, such that RP levels would select or filter for the translation of certain transcripts (6–8). The presence or absence of a RP on the ribosome also could allosterically interfere with conformational changes or alter interactions with ribosome-associated factors to change mRNA selection. RP-mediated selection of mRNAs could occur early in the initiation phase, by directly affecting ribosome recruitment, or otherwise alter the translation efficiency of specific transcripts at later steps. Our laboratory has previously utilized two RPs linked to such direct translation control, RPS25 and RACK1, to engineer human ribosomes for biophysical measurements (9, 10). These proteins are non-essential for ribosomal RNA (rRNA) maturation and proximal to ribosome-bound viral RNAs in cryo-EM-based models (**Figure 1A** and **Figure S1**) (11–13). Henceforth we use the term eS25 (by the modern RP nomenclature (14)) to describe the protein product of the human RPS25 gene, while the RACK1 protein and gene names are the same. Here we explore the biochemical and cellular basis by which these two RPs influence translational control.

**Figure 1.**
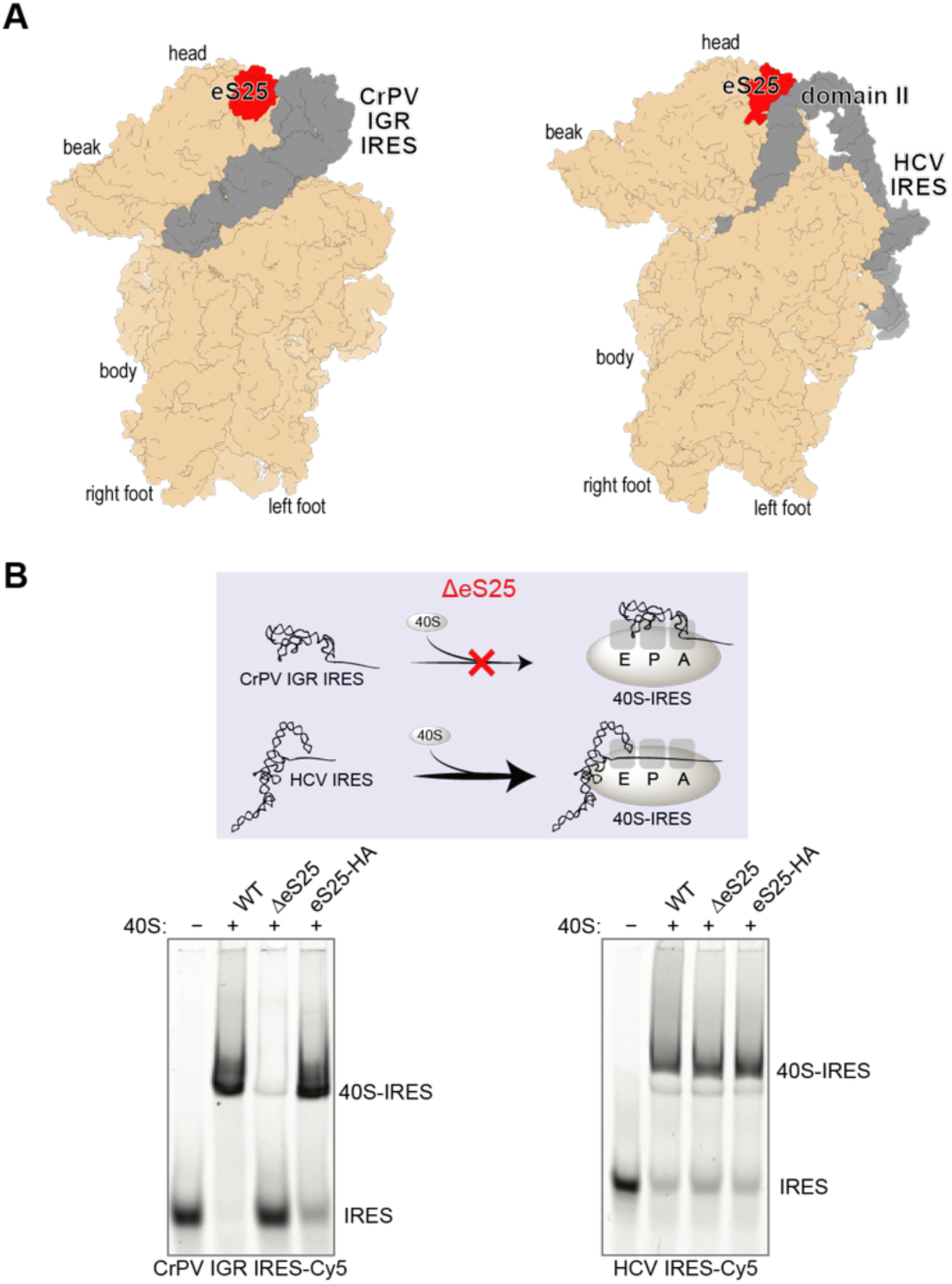
eS25 is not generally needed for direct 40S recruitment to internal ribosome entry sites. **A.** Structural models of the Cricket Paralysis Virus Intergenic Region internal ribosome entry site (CrPV IGR IRES, PDB 4v92, left) and Hepatitis C virus IRES (HCV IRES, PDB 5a2q, right). **B.** Native gel electrophoresis of WT, ΔeS25, and eS25-HA 40S ribosomal subunits binding to fluorescently labeled CrPV IGR IRES (left) or HCV IRES (right). Binding reactions were carried out with 30 nM labeled RNAs and 60 nM indicated 40S ribosomal subunits and resolved on acrylamide-agarose composite gels

eS25 is the archetype for RP-mediated translational control, as it is the sole RP for which *in vitro* and *in vivo* experiments converge to support a ribosome filtering model. Yeast ribosomes lacking eS25 have reduced affinity *in vitro* for the <u>C</u>ricket <u>p</u>aralysis <u>v</u>irus inter<u>g</u>enomic region internal ribosome <u>e</u>ntry <u>s</u>ite (CrPV IGR IRES) RNA, and the cells from which these ribosomes are isolated have reduced activity on a translational reporter *in vivo* (15, 16). These results are explained by structural analyses of CrPV IGR IRES-ribosome complexes, where eS25 forms direct interactions with the viral RNA near the E-site region of the ribosome (17–19) (**Figure 1A**, left panel). eS25 also directly contacts the Hepatitis C virus (HCV) IRES RNA within ribosome-IRES complexes (**Figure 1A**, right panel), and reporter assays concluded that eS25 is required for the HCV IRES to function (15). eS25 has been linked to the mechanism of other IRESs and to other specialized translation initiation events (20–22), and the observation that eS25 is sub-stoichiometric in cellular ribosomes has prompted the suggestion that eS25-mediated translational control is physiologically relevant (4).

Receptor for Activated C kinase 1 (RACK1) is also implicated in diverse translational and cell signalling processes, being first isolated based on interactions with protein kinase C and subsequent studies defined it as a ribosome “scaffold” for signalling proteins (23). RACK1 has been linked to translational processes including ribosome-associated quality control, reading frame maintenance, and IRES-mediated translation (24–27). The effects of RACK1 and RPS25 on signalling and translation have been mainly inferred from depletion experiments using siRNA knockdowns or genetic knockouts in yeast or human cells. Like eS25, reduction in cellular levels of RACK1 interferes with HCV IRES-mediated translation (26), but RACK1 does not form direct interactions with IRES RNAs on the ribosome or control ribosome recruitment *in vitro* (10).

The targeted disruption of cellular RP levels is the most accessible and rapid technique to examine RP function, yet such techniques come with limitations. Most critically, these assays cannot distinguish direct versus indirect effects. Both partial knockdown or full genetic knockout may cause immediate, direct effects on translation as well as long-term, indirect consequences on cell biology. Immediate effects might be obscured by the long lifetime of the ribosome (half-life of 5-7 days) and/or coordinated changes in overall ribosome levels in response to the loss of single RPs (28–30). Certain direct effects may be preserved in genetic knockouts, but they may also become muted due to cellular adaptation. The potential mechanisms for adaptation to RP loss are vast and might include the activated expression of a paralog gene, changes to ribosome assembly pathways or altered chaperone levels and degradation pathways to correct for challenges to protein homeostasis. Alterations in ribosome biogenesis may lead to the p53 stabilization by orphan RPs and MDM2 in mammals (31), but as shown by studies in yeast (which lack p53), may arise through other sources (32). The adaptation itself, rather than the RP’s normal function, could foreseeably lead to phenotypes that arise in genetic screens and reporter assays. Additionally, though acute knockdown experiments may be timed to minimize indirect effects from cellular adaptation, some effects may be unavoidable, and genetic rescue experiments have typically not been performed.

Here, we describe a series of experiments characterizing the direct and indirect effects that follow RP loss *in vitro* and in a haploid human cell line. With the aim of analysing direct effects, we first performed biochemical and compositional analysis of ribosomes isolated from RPS25 or RACK1 knockout cells. Prompted by the observation of an independent ribosome remodelling event, we turned our focus towards cellular analysis. We found that loss of RPS25 or RACK1 both drive indirect effects and partially overlapping, yet distinct sets of phenotypic outcomes. Given the centrality of eS25 to models of RP-mediated translational control, we scrutinized the RPS25 knockout cells and found that they had transitioned to a new state that drives several phenotypes previously assumed to be translational. Our findings uncover a host of indirect effects that accompany RP loss in a cell and have implications for the mechanistic interpretation of genetic lesions in the translational apparatus.

## RESULTS

### eS25 is not required for ribosome recruitment to the HCV IRES

We first sought to recapitulate in a human system the prior results from studies in yeast that demonstrated the *in vitro* requirement of eS25 for efficient ribosome recruitment to the CrPV IGR IRES (15, 16). We purified human 40S and 60S ribosomal subunits from wild type (WT), RPS25 knockout (KO), and RPS25-HA addback (AB) HAP1 cells (9), and *in vitro* transcribed and labelled CrPV IRES and HCV IRES RNAs (10). eS25-HA was present in polysomes and purified ribosomal subunits from the addback cell line, indicating successful ribosome incorporation (**Figure S1A-B**).

To determine the relative affinities of the CrPV IRES to the different 40S subunits, we used native gel analysis. Human eS25-deficient (ΔeS25) ribosomes showed decreased affinity to the IRES, as predicted by the yeast study (15), and affinity was rescue by eS25-HA addback 40S subunits (**Figure 1B** and **Figure S1C**). In contrast, the HCV IRES, which was not previously tested, showed no dependency on eS25 for 40S ribosome recruitment or magnesium-driven 80S complex formation (10, 33). These experiments demonstrate that the direct role of eS25 on ribosome recruitment by the CrPV IGR IRES is valid yet not a general rule of IRES-mediated translation, as currently thought (34). We reasoned that rather than through ribosome recruitment to the HCV IRES, eS25 and RACK1 might directly influence IRES-mediated translation at later steps of initiation or that the previous conclusions were due to indirect effects. However, the detailed biochemical nature of these RP knockout ribosomes was unclear and addressed below.

### RPS25 loss leads to partial remodelling of the large ribosomal subunit at eL22L1

To assess the composition of our purified ribosomes prior to pursuing additional biochemistry, we performed mass spectrometry on ribosomal subunits. We first examined the ribosomal subunits used above, which were purified in tandem with a WT control, and compared samples by LFQ intensities. RPs were readily detected in ribosomal samples, and we observed the expected reduction of eS25 and RACK1 levels in 40S subunits isolated from respective knockout cell lines (**Figure 2A** and **Figure S2**). Unexpectedly, upon similar analysis of purified 60S subunits, we detected an increase in the RP paralog eL22L1 in subunits from RPS25 knockout but not RACK1 knockout cell lines (**Figure 2A** and **Figure S2E**). We observed an increase in uL6 in the RPS25 knockout sample on this occasion, but found this was not reproducible, whereas we observed elevated eL22L1 in the ribosome samples of an RPS25 knockout on multiple occasions (**Figure S3-S4**).

**Figure 2.**
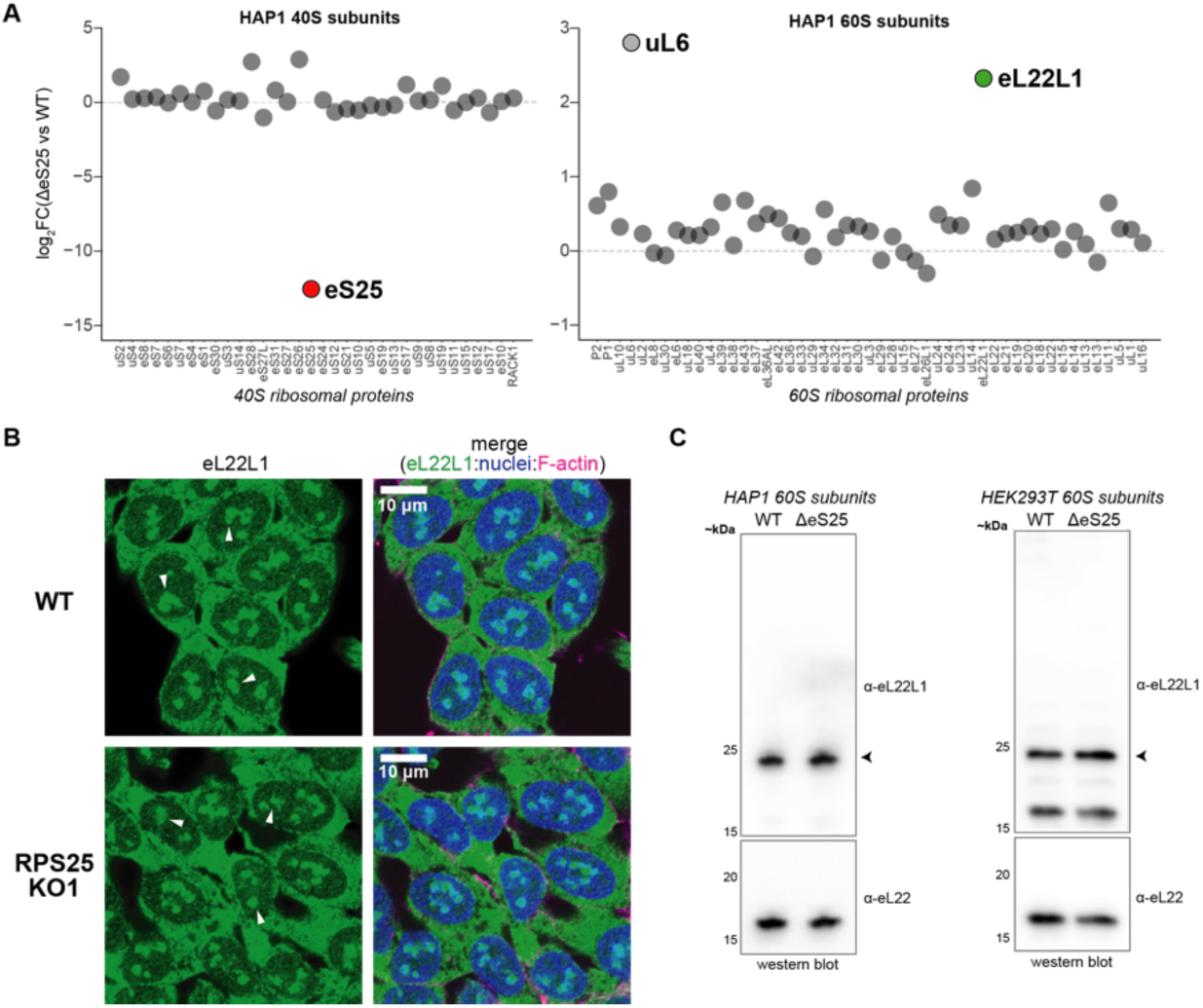
Genetic manipulation of RPS25 is imprecise at the ribosome level. **A.** Plot of the log2 fold-change (log_2_FC) in LFQ intensities between ΔeS25 and WT ribosomal subunits. **B.** Confocal images of fixed and stained WT and RPS25 KO1 HAP1 cells. Cells were stained with an antibody against eL22L1 alongside staining of nuclei with Hoescht and F-actin with Phalloidin 660. White arrows point to likely nucleolar structures. **C.** Western blot analysis of HAP1 and HEK293T 60S subunits for eL22L1. Arrows points to the major antibody-sensitive band, which likely corresponds to the long eL22L1 isoform.

In mice, RPL22 knockout leads to a compensatory upregulation of eL22L1 (35), and RPL22L1 has been implicated in extraribosomal and subnuclear roles in other settings (36–39). While most RP paralogs are very similar at the sequence level, human eL22 and eL22L1 are quite divergent, and depending on the isoforms being compared are ∼50-70% identical (**Figure S5**). Given the proposed subnuclear role and its presence on purified ribosomal subunits, we examined WT and RPS25 knockout cell lines by immunofluorescent (IF) imaging for eL22L1 (**Figure 2B**). eL22L1 was detected throughout the cytoplasm as well as subnuclear regions (likely nucleoli) (**Figure 2B** and **Figure S6**). To confirm the specificity of this antibody, we examined HAP1 60S subunits by immunoblotting and detected a product in both WT and KO ribosomes that is consistent with the expected molecular weight of the long isoform (∼21 kDa) (**Figure 2C**).

To probe how widespread the eL22L1 alteration is as a response to RPS25 loss, we analysed ribosomes from a couple other conditions. We first examined ribosomal subunits from an N-terminal mutant of RPS25 that was isolated in an attempt at making a knockout in HEK293T cells. As anticipated, eL22L1 was upregulated in ribosomes from this cell line by both mass spectrometry and immunoblotting (**Figure 2C** and **Figure S4**). Depletion of eS25 in the K562 cell line has been shown to mediate ricin toxin resistance (40), and in RNA-seq and ribo-seq analysis of these cell lines RPL22L1 was significantly elevated (G Hess, M Bassik, and N Ingolia, personal communication). To confirm the upregulation in this setting, we analysed 80S ribosomes from K562 cell lines harbouring shRNA targeting RPS25 or a non-targeting shRNA. We found that eL22L1 was again elevated in the RPS25-targeting samples by both mass spectrometry and immunoblotting (**Figure 2C** and **Figure S3**). These observations indicate that upregulated eL22L1 is a common response to RPS25 loss in human cancer cell lines.

Altered ribosome composition was recently described as a stress response in budding yeast and eL22L1 is upregulated in transformed human cell lines (**Figure S5B**) (41). eL22L1 expression also arises as a result of a mutation in an UFMylation pathway which primarily targets uL24 (RPL26) (42). eL22L1 therefore not only compensates for the loss of its own paralog, but it is a common adaptation occurring in settings where ribosome homeostasis and/or cellular identity has been disturbed. These observations indicate that genetic manipulations of RP genes do not precisely alter the ribosome composition, suggesting that indirect effects can even drive the remodelling of distal parts of the ribosome. We therefore halted our *in vitro* work and hypothesized that a stress response of the RPS25 knockout cells, rather than direct effects on translation, might explain the many phenotypes linked to RPS25.

### The HCV IRES does not require eS25 or RACK1 for its activity

eS25 and RACK1 are considered host factors for several viruses, and thought to mediate translation by IRESs and other non-canonical mechanisms (34). In addition to the CrPV IGR IRES, eS25 is considered essential for HCV IRES-mediated translation (15). To demonstrate this essentiality, Landry *et al.* depleted eS25 by siRNA knockdown in HeLa cells and measured HCV IRES-driven translation of firefly luciferase relative to that of cap-dependent Renilla luciferase from a dicistronic reporter construct, where a reduction of ∼50-75% IRES activity was observed (15). Similarly, knockdown of RACK1 was shown to reduce HCV IRES translation by ∼50% using mono-luciferase reporters in Huh7.5.1 cells (26).

To test whether these conclusions hold in knockout cells, we performed similar studies using HAP1 cell lines. In initial experiments, we observed large day-to-day variations in reporter activity and an altered growth behaviour of the knockout cell line. Since the HCV IRES dicistronic reporter activity has been reported to fluctuate by cell cycle (43), we designed an experiment to control for such variation (**Figure 3A**). We utilized two independent RPS25 KO clones—one isolated following CRISPR/Cas9 mutagenesis (KO1) and another following gene trap insertional mutagenesis (KO2), in addition to a RACK1 KO (9, 44, 45). While we observed a reduction of the relative luciferase signal in all mutants on this occasion, the relative luciferase level varied by cell density and is abolished in cells that have been synchronized by serum starvation (**Figure 3B-D** and **Figure S7A**). These results indicate that eS25 and RACK1 are not essential for HCV IRES-mediated translation and suggest another revision of the initiation factor requirements for the HCV IRES (46, 47). Reduction of HCV infectivity in eS25- and RACK1-depleted cells could instead result from indirect effects (15, 26). For RACK1 this notion is supported by other reports (48, 49), and we note that neither of these RPs emerged in genome-wide screens for resistance to HCV infection and replication (45, 50, 51).

**Figure 3.**
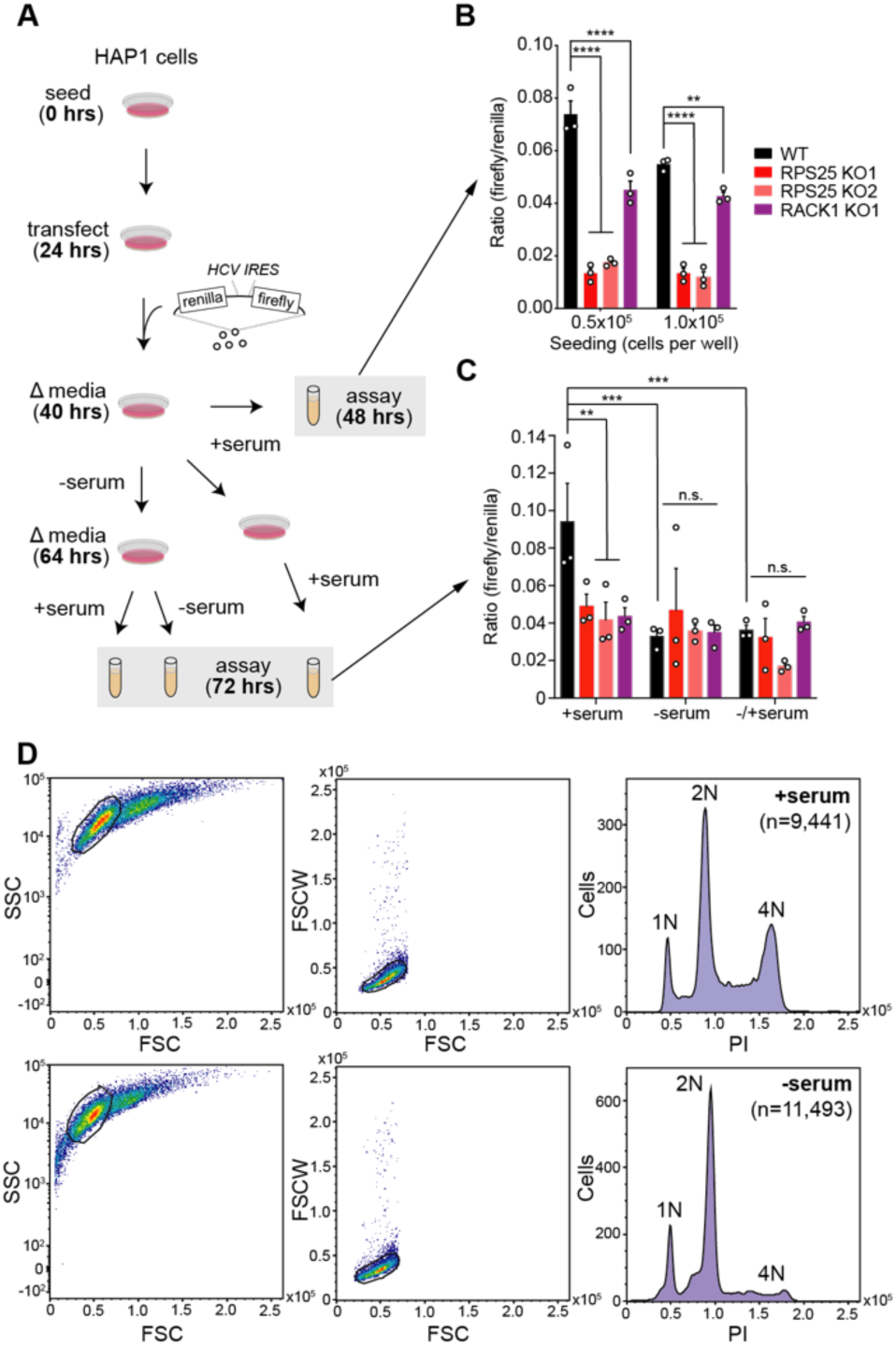
RPS25 is not required for HCV IRES-mediated translation. **A.** Experimental scheme for cell cycle synchronization by serum-starvation and subsequent dual-luciferase assays. **B.** Results from assays of WT and KO cells at 24 hours post-transfection at two seeding densities. **C.** Results from assays of WT and KO cells at 48 hours post-transfection at the low seeding density (0.5×10^5^ cells/well), under different serum conditions. Error bars in B-C represent the SEM from three biological replicates. The cell lines and conditions were analyzed by two-way ANOVA and a Fisher’s LSD test was used to determine statistical significance, without taking into account multiple comparisons. This exact experiment was performed on a single occasion, while individual perturbations using the same reporter were performed on separate occasions with similar results. P-values: ≥ 0.05 (n.s.), 0.001-0.01 (**), 0.0001-0.001 (***), and <0.0001 (****). **D.** Example propidium iodide (PI) FACS-based assay to verify cell cycle arrest under serum-starvation conditions. Single cells were identified by manually drawing a window on a plot of Side Scatter (SSC) versus Forward Scatter height (FSC) (left panels), followed by on a plot of Forward Scatter Width (FSCW) versus FSC. Histograms of the single cell intensity from the BluFL2 PMT are shown.

### RPS25 is not required for flavivirus translation

RPS25 is a prominent hit in genome-wide haploid genetic screens for flavivirus host factors, including dengue virus (DENV) and the closely-related Zika virus (ZIKV) (45, 52). siRNA studies in other cell lines have further supported the role of RPS25 and RACK1 as host factors for DENV and other flaviviruses and suggested that this effect is due to translational control (53, 54). Flaviviruses are positive-sense RNA viruses and have capped genomic RNAs (gRNAs) implying that their translation could share a similar mechanism with canonical cellular capped mRNAs. Given the connection to IRES-mediated translation events, it has been interpreted that the DENV resistance phenotype of RPS25-deficient cells is due to a specialized translation mechanism such as the recent suggestion that the DENV and ZIKV 5’ untranslated regions harbour certain IRES properties (53, 55).

To test the reported flavivirus resistance phenotype of the RPS25 knockout, we employed a DENV infectious clone encoding a Renilla luciferase sequence (DENV-luc) (45). DENV infection kinetics enable the dissection of its lifecycle: in the first ∼8 hours following infection, DENV enters cells and translates its genome, and only after this time does the replication phase become dominant (56). We therefore performed a time-course infection experiment with DENV-luc and WT, RPS25 KO, and addback cells. While a strong reduction of luciferase was observed at 24 hours post-infection for the RPS25 KOs, we observed only minor differences between cell lines at 4- and 8-hours post-infection, indicating that eS25 is not required for viral entry or initial translation (**Figure 4A-B** and **Figure S8A**). Unexpectedly, the eS25-HA addback failed to rescue the viral attenuation at 24 hours in either knockout cell line, despite our previous observation that eS25-HA incorporates into ribosomes and recovers *in vitro* ribosome recruitment to the CrPV IRES (**Figure 1B** and **Figure S1A-C**).

**Figure 4.**
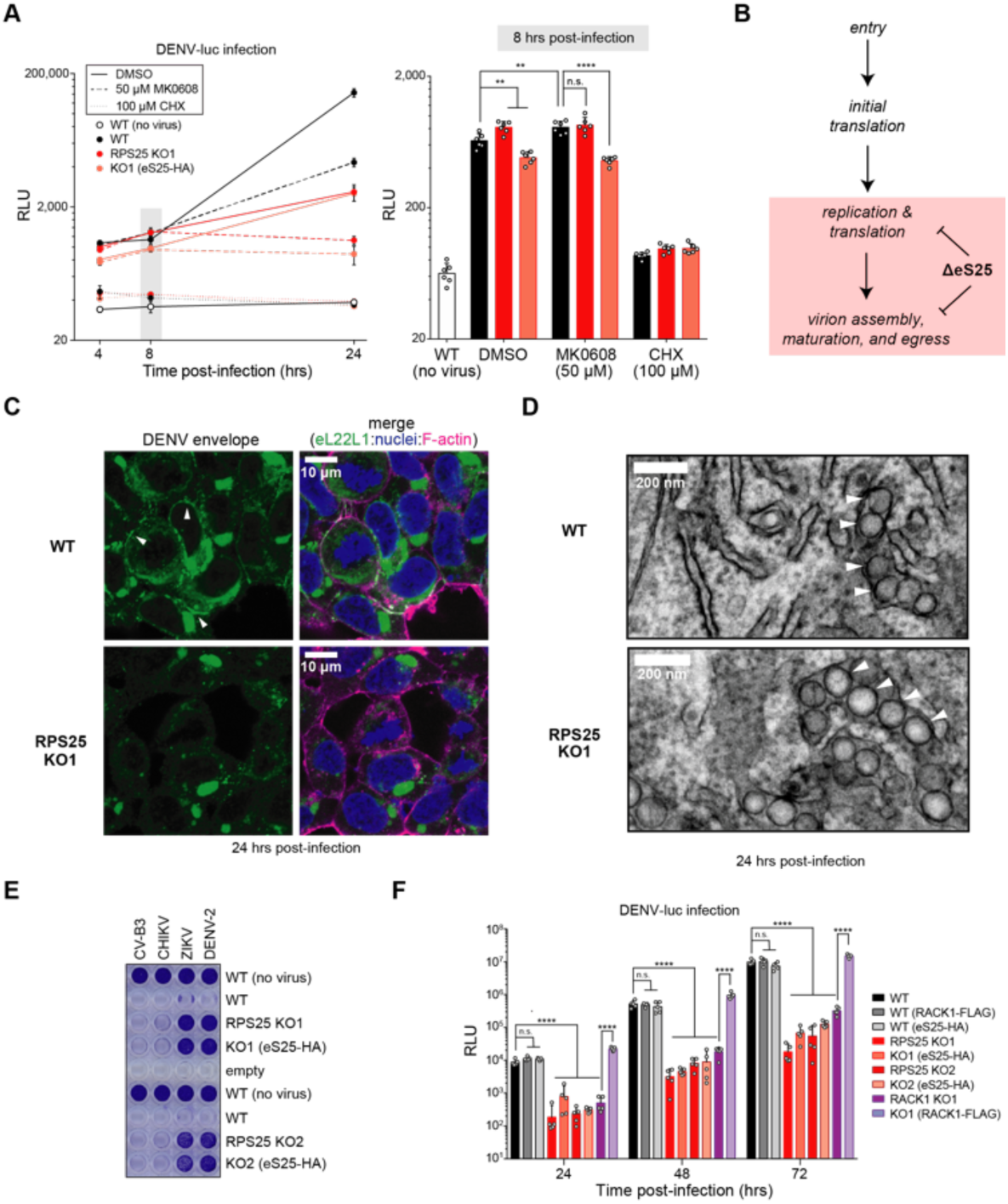
RPS25 loss indirectly inhibits flavivirus infection. **A.** DENV-luc infection in WT, RPS25 KO, and eS25-HA AB cells. At time 0, cells were infected with MOI=0.05 DENV-luc and assayed 4, 8, and 24 hours post-infection with drug treatments to inhibit viral replication (MK0608) or mRNA translation (CHX). Renilla luciferase activity is measured in relative light units (RLUs). Left panel shows the averages of n=6 biological replicates for each cell line and time point with error bars representing the 95% CI, while right panel shows the full data for the 8-hour time point. Statistical significance was determined with a two-way ANOVA and Tukey multiple comparison test for each cell line and treatment at respective time points. Only comparisons for DMSO- and MK0608-treated cells are shown. A similar result was observed from two independent experiments. **B.** Model for the defect in DENV infection following RPS25 loss in HAP1 cells, wherein the major effect appears >8 hours post-infection when translation and replication are coupled. **C.** Confocal images from staining of WT and RPS25 KO HAP1 cells infected with DENV-2 at MOI=2. Fixed cells were stained with an antibody against the dengue envelope (E) protein alongside staining of nuclei with Hoescht and F-actin with Phalloidin 660. White arrows point to bead-like structures of the E protein within the cell periphery of infected WT cells. **D.** Transmission electron microscopy (TEM) images of an RPS25 KO cell section prepared following infections with DENV-2 at MOI=2. White arrows indicated ER membrane-derived replication vesicles. Full images for C-D are in Figure S8. **E.** The cell killing effect of four viruses was assayed with crystal violet assays. Cell lines were infected with the Coxsackie B3 virus (CV-B3, MOI=1), Chikungunya virus (CHIKV, MOI=1), Zika virus (ZIKV, MOI=25), and DENV serotype 2 (DENV-2, MOI=25). Similar results were observed in three independent experiments. **F.** Cells were infected with DENV-luc (MOI=0.018) and assayed at 24, 48, and 72 hours post-infection. Error bars and statistical significance as in (A) with n=5 biological replicates. P-values: ≥ 0.05 (n.s.), 0.001-0.01 (**), and <0.0001 (****).

To understand the nature of flavivirus resistance at late stages of infection, we imaged DENV-infected WT and RPS25 KO cells stained for structural (envelope, E) and non-structural (NS3) proteins. In all cases, DENV proteins were specifically detected in infected WT and knockout cells (**Figure 4C** and **Figure S8-11**). Staining for both viral proteins was most prominent adjacent to nuclei, likely corresponding to the ER, and we observed E protein staining at the cell boundary, likely corresponding to budding virions (**Figure 4C**). The levels of DENV proteins were visually attenuated in both RPS25 knockouts versus WT, yet E protein staining was largely absent at the cell boundary in both knockouts (**Figure 4C** and **Figure S8C**). This finding raised the possibility that ultrastructural changes might be present in the RPS25 knockout in the context of DENV infection, such as alterations in the ER membrane-derived replication vesicles (57). We therefore analysed WT and RPS25 knockout cell slices by transmission electron microscopy (TEM) following 24 hours of DENV infection. We readily observed DENV-induced vesicles in both knockout clones that were absent in uninfected cells (**Figure 4D, Figure S8**, and **Figure S12**). We also observed virions, both at the ER and at the cell periphery (**Figure S8**). The DENV-induced replication vesicles were not qualitatively different in size and shape between WT and knockout cells, suggesting that RPS25 loss did not impair this crucial aspect of the lifecycle. These results therefore indicate that the flavivirus resistance of the RPS25 knockout derives from an indirect mechanism that arises at late stages of infection and is unrelated to translation.

To confirm that the eS25-HA addback fails to rescue DENV infection, we performed additional viral infection experiments. First, using a small panel of untagged viruses on HAP1 cell lines, we found that the protection against DENV- and ZIKV-mediated cell death was observed for both RPS25 knockouts, and specific for flaviruses as shown by the lack of protection against a picornavirus (Coxsackievirus B3, CV-B3) and an alphavirus (Chikungunya virus, CHIKV) (**Figure 4E**). Again, both addback clones failed to rescue the WT sensitivity to DENV or ZIKV. We confirmed this resistance phenotype in both RPS25 knockouts via plaque assays and immunoblotting structural and non-structural DENV proteins (**Figure S13A**). Finally, performing DENV-luc infections on additional cell lines for multiple durations, we found that the protection against DENV-luc in both RPS25 knockouts was clear and significant across the first 72 hours of infection, and the eS25-HA addback consistently failed to rescue sensitivity (**Figure 4F** and **Figure S13B**). In contrast, we observed a reduction in DENV-luc signal in two independent RACK1 knockout clones and found that expression of RACK1 cDNA (RACK1-FLAG) rescued this defect. Expression of RPS25-HA or RACK1-FLAG in WT cells did not noticeably alter DENV-luc signal, arguing against a dominant-negative effect from the tagged proteins. These results therefore demonstrate that viral resistance phenotype from RPS25 knockout is robust, flavivirus-specific, and cannot be rescued by functional cDNA expression.

### RPS25 loss drives diverse phenotypes that cannot be rescued

RPS25 scores as a negative regulator in several genetic screens exploring unrelated phenotypes (**Figure S14A**) (58). To understand the phenotypic diversity of the RPS25 knockout and whether phenotypes can be rescued in other settings, we intersected the results from a DENV resistance screen with that from a spliced Xbp1 (Xbp1s) screen (**Figure 5A**) (45, 58). Perhaps unsurprising given their roles in endoplasmic reticulum (ER) homeostasis, several of the ER-resident DENV host factors also increase cellular levels of spliced Xbp1 in response to an 8-hour treatment with 2 μM tunicamycin (Tm) (**Figure 5A**) (58). However, the appearance of RPS25 in the Xbp1s screen as the only RP was surprising given that it is not directly related to ER function.

**Figure 5.**
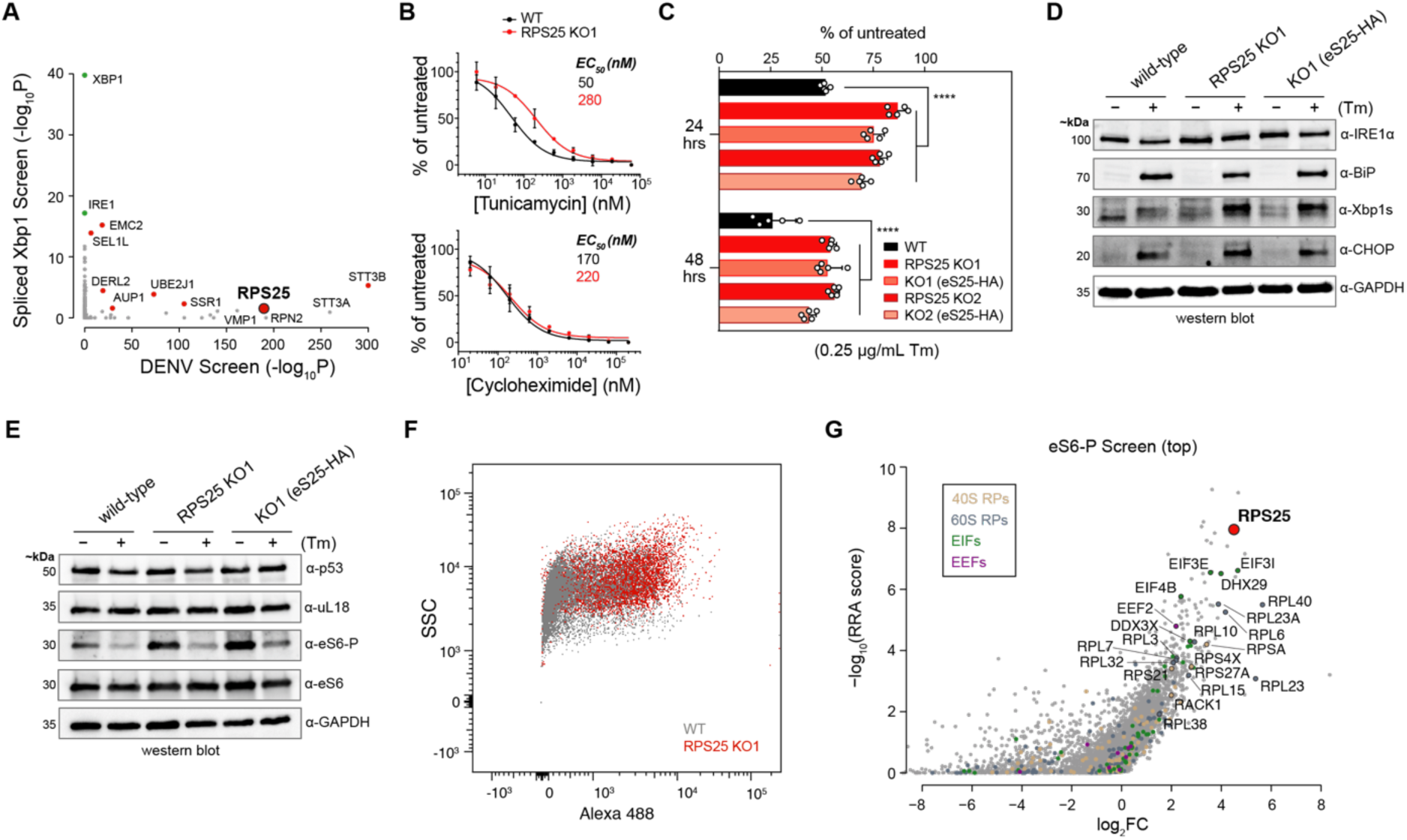
RPS25 loss drives pleiotropic phenotypes that cannot be rescued. **A.** Intersection of published genetic screen results depicts shared regulators of DENV resistance and Xbp1 splicing in HAP1 cells. The -log10 FDR-corrected P-value (−log_10_P) of the spliced Xbp1 screen (58) was plotted against that of the DENV screen (45). The Xbp1 screen was FACS-based and identified both positive and negative regulators of the pathway, while the DENV screen was a live/dead screen and all mutations therefore promote resistance to DENV-mediated cell death. Green dots represent expected positive regulators, and red dots represent shared hits between the two screens. **B.** RPS25 KO cells are resistant to Tm-mediated cell death. Cells were assayed by an MTT assay 48 hours after drug treatment and the absorbance at 570 nm was normalized to untreated cells. **C.** Resistance to Tm-mediated cell death is a common and irreversible phenotype from RPS25 loss in HAP1 cells. As in (B), HAP1 cells were assayed by an MTT assay, but using a single Tm concentration (0.25 μg/mL). For B-C, error bars represent the 95% CI of n=5 biological replicates. Similar results were observed in two independent experiments. P-values: ≥ 0.05 (n.s.) and <0.0001 (****). **D-E.** Western blots of cell lysates from WT, RPS25 KO, and eS25-HA AB cells treated with or without a low dose of Tm (0.25 μg/mL) for 24 hours. The full blot for (E) is in Figure S14. **F.** Flow cytometry plot of singlets from fixed WT and RPS25 KO cells stained with primary anti-eS6-P antibody and secondary Alexa 488 antibody. Side scatter (SSC) is plotted against the antibody staining fluorescence at 525nm (B525, Alexa 488). The number of cells (n) are 84,596 and 65,182 for WT and KO cells, respectively. **G.** Positive selection gene scores (−log_10_(RRA score)) versus fold-change (log_2_FC) of CRISPR-Cas9 guides in the high eS6-P population (top) from genome-wide screen of eHAP cells.

We reasoned that appearance of RPS25 as a top hit in a ricin toxicity screen (40), and the utilization of ERAD components for ricin toxin processing, might suggest that altered protein homeostasis and secretory pathway dysfunction is a common indirect effect of RPS25 loss. We therefore examined the ER stress phenotypes in HAP1 cells as a window into this relationship. We titrated tunicamycin onto WT and RPS25 KO1 cells, and measured EC_50_ values for growth inhibition using proliferation assays. We observed an increase in the EC_50_ value for the RPS25 knockout versus WT cells after 48 hours of treatment (**Figure 5B**). In contrast, parallel titration of the translation inhibitor cycloheximide (CHX) onto WT and RPS25 knockout cells did not differentially affect the cell lines, indicating that toxin-resistance is not general. To test the robustness and reversibility of the tunicamycin resistance, we treated both RPS25 knockout clones and their respective addbacks with a low dose of tunicamycin for 24 and 48 hours and assessed resistance with a proliferation assay. As with the flavivirus resistance phenotype, we found that the effect is significant for both knockout clones and not rescued by eS25-HA (**Figure 5C**).

Tunicamycin resistance is a general response to RP knockout in yeast, where this had been generally attributed to reduced ER burden through diminished protein synthesis (1). To understand the resistance of the RPS25 knockout, we measured UPR markers in response to drug treatment by RT-qPCR (59). We found that tunicamycin activated Xbp1 splicing and increased ATF4 levels in the RPS25 knockout similar to WT (**Figure S14C**). Further, the low dose tunicamycin treatment increases levels of Xbp1s in the RPS25 knockout, which were not rescued by the addback (**Figure 5D**). Since sustained tunicamycin treatment in yeast drives aneuploidy-enabled resistance (60), we karyotyped the RPS25 knockout clones to ensure that no chromosomal alterations had taken place (**Figure S14D**). We observed no evidence of aneuploidy besides the expected abnormalities of the parental HAP1 clone (61), indicating that the many stable alterations cannot be explained by such a mechanism. Together, these findings indicate that RPS25 loss drives a potent ER stress resistance phenotype, which like flavivirus resistance, cannot be rescued by eS25-HA.

We hypothesized that RPS25 loss drives many indirect alterations to cellular protein homeostasis, of which flavivirus and ER toxin resistance are only two. To assess a broader catalogue of proteins related to cellular and ribosome homeostasis, we probed cell lysates with a panel of antibodies (**Figure 5E** and **Figure S14E**). We observed no clear alterations of small or large subunit RPs or the tumour suppressor p53, indicating that neither ribosome levels or an ongoing p53 stress response can explain these alterations. Most notably, eS6 phosphorylation (eS6-P) was elevated in the RPS25 knockout and addback relative to WT, suggesting alterations to mTOR signalling and autophagy (62, 63). To assess the specificity and robustness of this effect, we examined lysates from both RPS25 knockout clones, a RACK1 knockout, and all respective addbacks. We found that the stable elevation in eS6-P in the RPS25 knockouts was reproducible, and eS6-P was also elevated in the RACK1 knockout but rescued by its addback (**Figure S14F**). We observed a similar trend with the autophagy biomarker LC3B, where both RPS25 knockouts have an irreversible increase in LC3B-II, which is also elevated in the RACK1 knockout but rescued by its addback (**Figure S14F**). RPS25 loss therefore appears to induce autophagy similar to other RP gene mutations (62), and like all cellular phenotypes we tested this cannot be rescued by the addback.

The strong elevation in phosphorylated eS6 in both RPS25 and RACK1 knockouts suggested that eS6-P may be a general response to ribosomal protein loss (62). To test this possibility, we leveraged wild-type and RPS25 knockout cells to optimize immunoflow conditions for a genome-wide CRISPR/Cas9 screen of eS6-P levels (**Figure 5F** and **Figure S15A**). We mutagenized the HAP1-derived eHAP cell line and sorted out the high and low populations of eS6-P stained cells for deep sequencing. Strikingly, the population with high eS6-P was enriched for many RP gene targeting guide RNAs, of which RPS25 was the most strongly enriched (**Figure 5G**). Guides for RPs and translation-related were broadly enriched in the high eS6-P population, but not the low population, and both populations yielded known components of the mTOR and RPS6K signalling pathways (**Figure S15B** and **Supplementary Table 4**). By defining the genetic regulators of eS6-P, this screen therefore reciprocally validated the strong induction of eS6 phosphorylation in the RPS25 knockout and further illustrated the imprecision of RP mutations genes on the cellular ribosome pool.

Given the many alterations of the RPS25 knockout that cannot be rescued and the suggestion that eS25 is required for efficient repeat-associated non-AUG (RAN) translation that occurs in neurodegenerative diseases (22), we asked whether such a phenotype can be rescued. Immunoblotting WT, RPS25 knockout, and addback cell lysates following transfection with a C9orf72 66-repeat (C9-66R) expression construct demonstrated a failure of the addback to rescue WT expression levels of the polyGA RAN translation product (**Figure S14G**). Since no clear differences in C9-66R mRNA stability were observed in the previous study (22), these findings suggest that post-translational mechanisms of dipeptide clearance (such as autophagy) may instead explain the strong reduction in polyGA in the HAP1 RPS25 knockout. Since acute reductions in eS25 were shown to alter dipeptide repeat levels in other settings, more work is needed to decipher whether there is a role for translation in those conditions. However, in the HAP1 cells used here and in the prior study, the strong reduction in polyGA accumulation cannot be rescued.

### RNA-seq and membrane mass spectrometry define a common cell state

We were perplexed by the failure of the RPS25 addback to rescue phenotypes and therefore hypothesized that knockout cells had transitioned to a new cell state, which itself drove the pleiotropy. To uncover the transcriptional basis of such a state, we applied RNA-seq to wild type, RPS25 knockouts, and addback HAP1 cells. As predicted by prior measurements, most RP transcripts were present at similar levels, while RPS25 transcripts were strongly reduced and RPL22L1 was elevated in the RPS25 knockouts (**Figure 6A** and **Figure S16A**). The most significantly upregulated gene in both conditions is ANXA1, encoding Annexin A1—an anti-inflammatory protein that participates in innate immunity (64).

**Figure 6.**
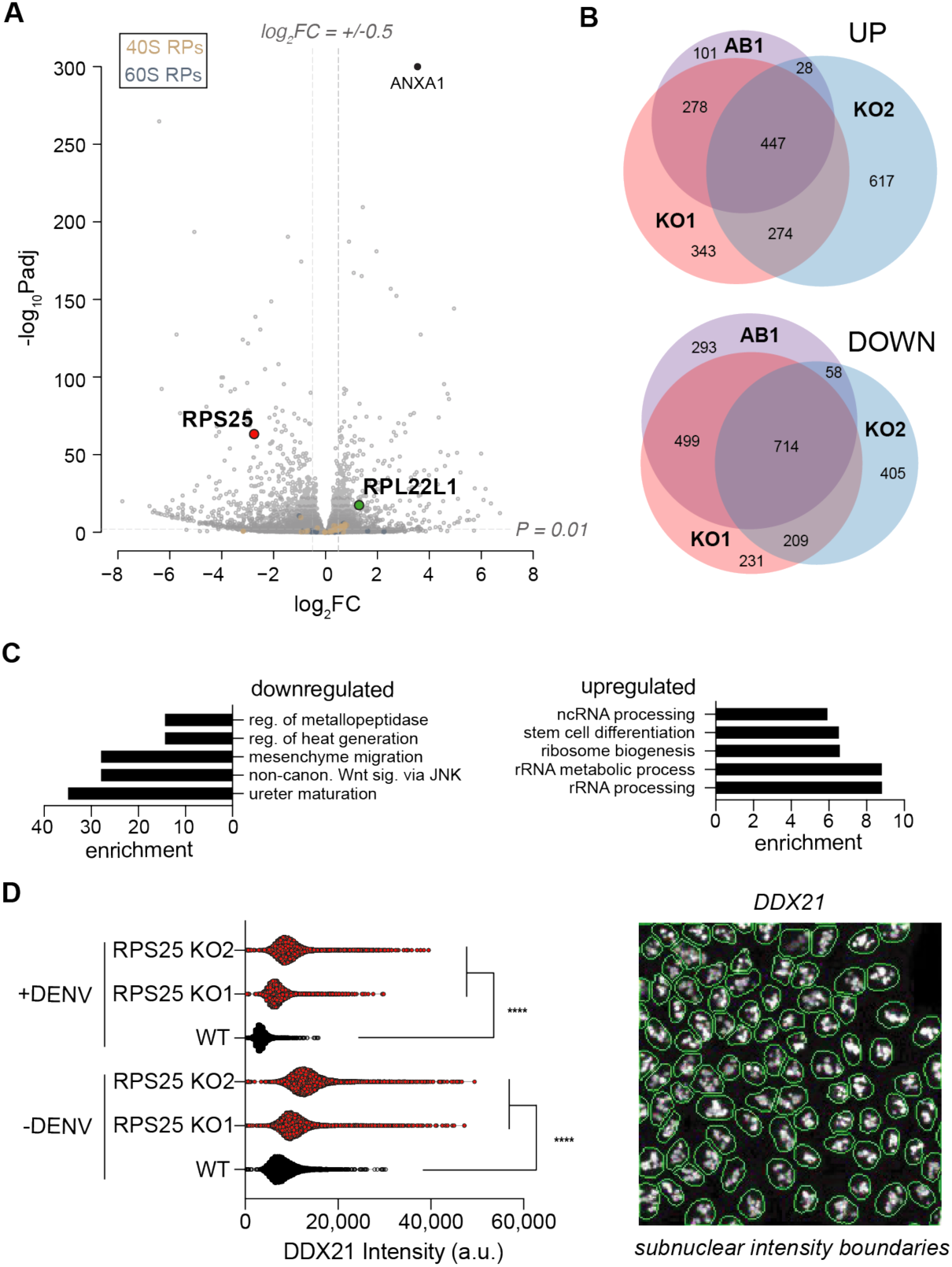
RPS25 loss drives a common dysregulation of ribosome biogenesis. **A.** Plot of RNA-seq fold-changes for RPS25 KO1 versus WT HAP1 cells (n=6). Y-axis represents the negative log10 FDR-adjusted P-values (−log_10_Padj), while x-axis represents the log2 fold-change (log_2_FC). To indicate cutoffs for ontology analysis, horizontal and vertical gray lines are shown. **B.** Venn diagrams of RNA-seq upregulated and downregulated genes between RPS25 KO1, RPS25-HA AB1, and RPS25 KO2 HAP1 cells. Gene lists are based on the significance and FC cutoff described in (A). **C.** GO analysis for Biological Processes from shared genes in the RPS25 KOs and ABs. Lists are based on the overlap of all four conditions as in in Figure S16, showing the top five significant GOs (FDR-corrected P-value<0.05) with their fold enrichment. **D.** Quantitative image analysis of WT and RPS25 KO for subnuclear DDX21 signal. Right viewfield represents an example of automated nuclear boundary identification for use in image analysis (full images in Figure S17). Violin plots depict intensity values from DDX21 antibody staining from within nuclear regions. The number of cells analyzed (n) for each condition are as follows: 7,229 (WT-DENV), 9,076 (KO1-DENV), 10,199 (KO2-DENV), 5,747 (WT+DENV), 8,405 (KO1+DENV), and 10,093 (KO2+DENV). Statistical significance represents the results of a two-way ANOVA for each cell line and condition, correcting for multiple comparisons with a Tukey test. P-value: <0.0001 (****).

We established fold-change cut-offs for differentially expressed genes and examined the overlap for upregulated and downregulated transcripts (**Figure 6A-B**). Despite certain clonal differences, both knockouts share common, irreversible phenotypes so we focused on shared changes from all conditions using gene ontology (GO) analyses. By analysis of cellular components, we observed the upregulation of GOs related to events in the nucleolus, and downregulation of GOs related to the extracellular matrix, cell communication, and the ER lumen (**Figure S16E**). Examination of upregulated biological process ontologies revealed a number of categories related to ribosome biogenesis, followed by stem cell differentiation (**Figure 6C**).

To understand these common changes, we clustered genes and visualized the results as heatmaps (**Figure S16F**). Nucleolus-related genes were broadly upregulated, but at modest levels. To validate this finding, we used quantitative analysis of IF images from staining the protein products of altered nucleolar protein-coding transcripts (DDX21 and eL22L1, **Figure 6D** and **Figure S17**). Both RPS25 knockouts have elevated DDX21 signal in subnuclear regions without notable changes in nuclear area (**Figure 6D**). With eL22L1, which also has a prominent extranuclear localization, such a trend was not clear (**Figure S17C**). Infection of cells with DENV attenuated the DDX21 signal within nuclei for all cell lines (**Figure 6D**), which may be explained by the finding that DENV proteins (NS5 and capsid) shuttle to the nucleolus and likely alter its function like other viruses (65–67). These data therefore demonstrate a common signature of nucleolar dysregulation elicited by RPS25 loss.

To reveal other transcriptional changes that cannot be rescued and therefore define the cell state, we examined select shared ontologies from the RNA-seq dataset. Stem cell differentiation was a top upregulated GO with strong, stable alterations, and mesenchyme migration and non-canonical WNT signalling ontologies were also altered (**Figure S16C**,**E**). Potentially consistent with the indication of ER stress resistance in the RPS25 knockouts, many ER lumen transcripts were also strongly and irreversibly altered (**Figure S16F**). The transcriptional analysis therefore demonstrates the dysregulation of differentiation-related genes and widespread alterations that cannot be rescued.

In order to validate prominent biomarkers of the state change, we performed mass spectrometry on ER membranes extracted from cells (**Figure S18A**) (68). The purification identified many known ER proteins and RPs at high LFQ intensities (**Figure S18C**), and comparison of significant fold-changes between knockout and WT membrane was well validated by the strong reduction in eS25 and increase in eL22L1 (**Figure 7A** and **Figure S18D**). To delineate common changes in both knockouts, we established cut-offs for differentially expressed proteins and identified sets of upregulated and downregulated proteins for ontology analysis (**Figure 7A** and **Figure S18F**). Cytosolic ribosome-related proteins were upregulated as was mitochondria gene expression, and ER-related processes, cellular biosynthesis and metabolism were all downregulated (**Figure S18G**). To establish the overlap with RNA-seq measurements, we intersected the differentially expressed genes from both experiments and identified shared genes (**Figure 7B**).

**Figure 7.**
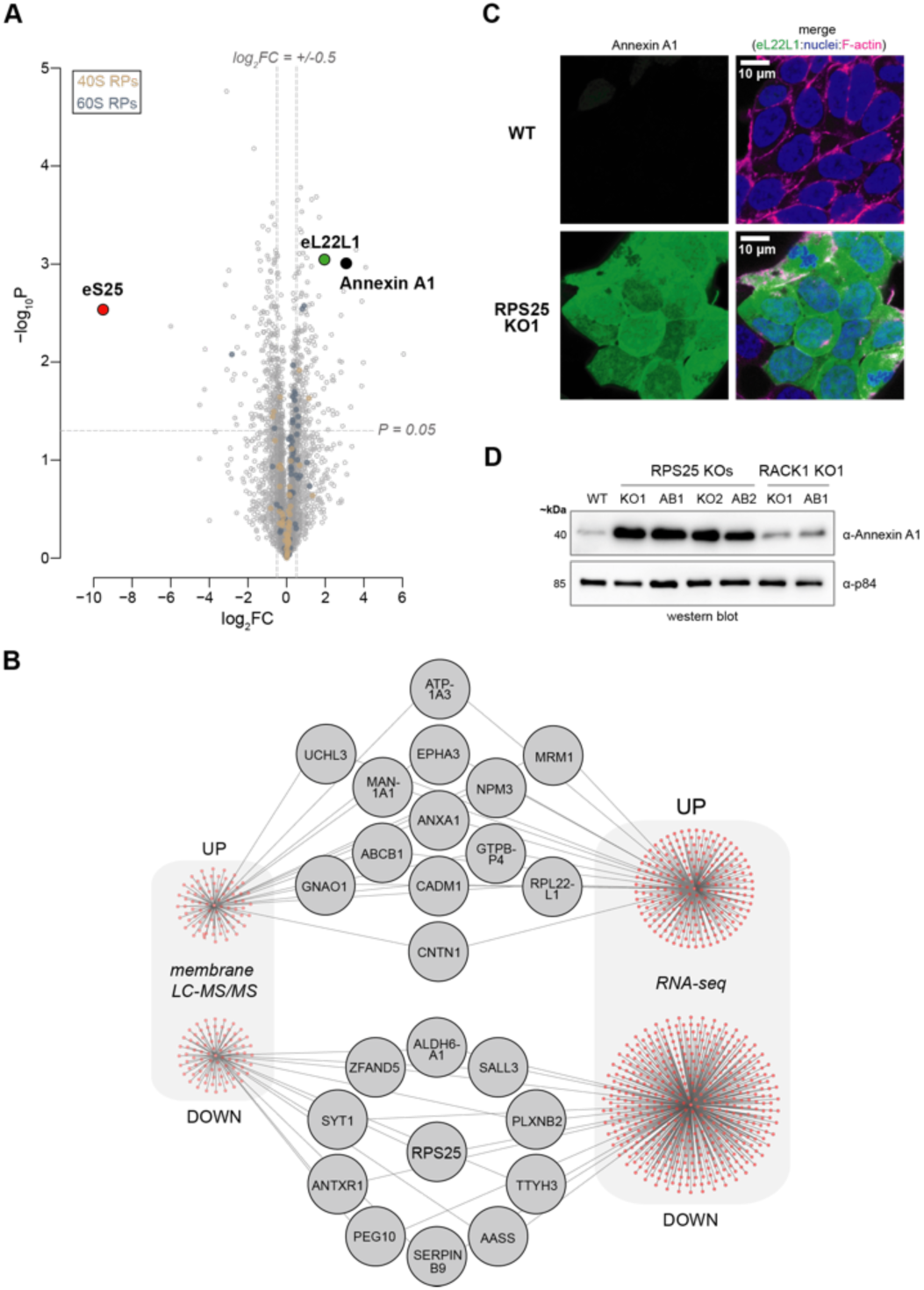
Membrane mass spectrometry confirms a common RPS25 knockout cell state. **A.** Plot for fold-changes from LFQ intensities based of RPS25 KO1 versus WT samples. Y-axis represents the negative log10 P-values (−log_10_P-value), while x-axis represents the log2 fold-change (log_2_FC). Values are derived from analysis of three biological replicates for each cell line and a two-tailed heteroscedastic t-test. To indicate cutoffs for ontology analysis, horizontal and vertical gray lines are shown. **B.** Network plot intersecting significant upregulated and downregulated genes shared between membrane LC-MS/MS and RNA-seq analyses. **C.** Confocal images from IF imaging of WT and RPS25 KO1 HAP1 cells for Annexin A1. Fixed cells were stained for Annexin A1, nuclei (Hoescht), and F-actin (Phalloidin 660). Full images in Figure S18. **D.** Immunoblot analysis of HAP1 whole cell lysate for Annexin A1 demonstrates specificity to the RPS25 KOs.

Given the prominence of ANXA1 in both data sets, we validated this change by microscopy and immunoblotting. By imaging, Annexin A1 was strongly expressed throughout the knockout but not WT cells (**Figure 7C** and **Figure S19)**. Elevated Annexin A1 levels could not be rescued by eS25-HA in the RPS25 knockouts, whereas Annexin A1 is at wild-type levels in the RACK1 knockout (**Figure 7D**), indicating that its upregulation is not a general response to ribosome stress or cellular fitness defects. Given the role of Annexin in innate immunity (64, 69), these findings suggest that RPS25 loss may have primed cells with heightened antiviral and antitoxin immunity for which elevated Annexin A1 is an participant or consequence.

### RPS25 loss-driven phenotypes cannot be corrected by genomic locus repair

In every cellular assay we examined, eS25-HA failed to rescue phenotypes. We felt confident that this was not due to altered function of the transgene since multiple experiments demonstrated successful expression, ribosome incorporation, and *in vitro* functionality of eS25-HA. Nevertheless, the protein tag could interfere with some but not other functions, the mRNA is expressed from a non-native promoter, and the mature mRNA is produced without splicing from pre-mRNA. To confirm the irreversibility of phenotypes and expression markers, we therefore repaired the mutated genomic locus in the knockout cell line. We designed CRISPR/Cas9 guides targeting regions of the RPS25 locus upstream and downstream of a 6-nt deletion in the RPS25 KO1 clone (**Figure S20A**,**C**), and applied homology-directed repair with CRISPR/Cas9 and a template of the wild-type sequence. We identified revertants by PCR-based screening following two repair strategies (HDR1 and HDR2), verified eS25 expression by immunoblotting, and confirmed the genomic repair by sequencing genomic DNA amplicons (**Figure 8A** and **Figure S20B**,**D**). We then asked how the HDR clones respond to DENV infection and whether the elevated expression of Annexin A1 marker was reversed. Consistent with a state change, all clones failed to rescue the sensitivity to DENV-luc infection and express Annexin A1 at levels similar to the knockout (**Figure 8B**). The knockout, addback, and HDR clones therefore appear to be phenotypically indistinguishable. As a result, we conclude that these and other knockout phenotypes not rescued by the eS25-HA addback are the indirect products of an irreversible cellular state change following RPS25 loss (**Figure 8C**).

**Figure 8.**
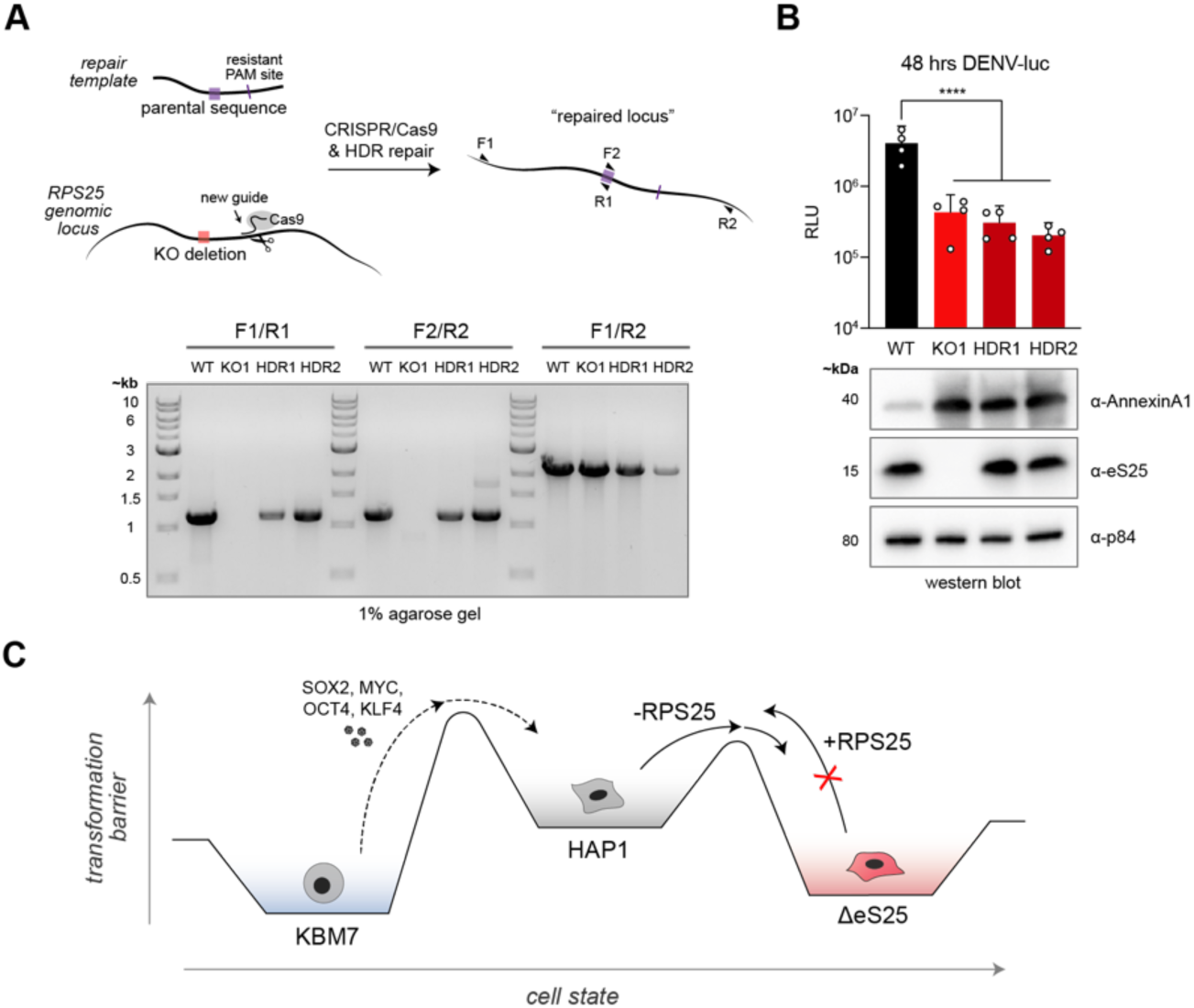
The phenotypic and expression state of RPS25 loss cannot be rescued by genomic repair. **A.** Homology-directed repair (HDR) of the RPS25 KO1 HAP1 cells line. Upper repair schematic depicts locus repair with a new guide targeting RPS25 intron 1. Agarose gel electrophoresis of PCR fragments from screening for repair clones HDR1 (2B11) and HDR2 (6G11). **B.** Immunoblot analysis and DENV-luc assay of WT, RPS25 KO, and RPS25 HDR repair clones. Top, luciferase values were assayed for each cell line at 48 hours post-infection with DENV-luc (MOI=0.014). Error bars represent the 95% confidence interval (CI) of n=4 biological replicates. Statistical significance represents the results of a one-way ANOVA, correcting for multiple comparisons with a Tukey test. Bottom, western blots of post-nuclear lysate demonstrate that high Annexin A1 expression is not reversed in repair clones. The same experiments were performed on a second occasion with similar results. P-value: <0.0001 (****). **C.** A model for the cellular transformation of the RPS25 KO cell line to explain the phenotypic hysteresis. The x-axis represents a non-dimensional cell state, while the y-axis represents a transformation barrier between cell states. The HAP1 cell line was previously derived by treating the near-haploid KBM7 cell line with the Yamanaka factors (3).

## DISCUSSION

Our results illustrate that genetic lesions of ribosomal protein genes are imprecise at the ribosomal and cellular level, and that RPs with suggested roles in direct translational control are no exception. We propose that RPS25 loss elicits a specific cellular state change, which itself drives phenotypes. Functional cDNA expression could not rescue any phenotype we tested, including flavivirus resistance, ER toxin resistance, and RAN translation, which were all previously assumed to result from specialized translation mechanisms. These irreversible phenotypes correlate with multiple stress-related expression markers and suggest that knockout cells have dysregulated autophagy—a mechanism which itself could foreseeably degrade DENV proteins and RAN translation products. Since RPS25 is associated with a unique set of phenotypes compared with RACK1 and other RP mutants, the irreversible state is unlikely to arise by random clonal divergence (**Figure S14A**). RPS25 knockouts made by different strategies behave similarly and have a common rewiring of gene expression, indicating that the state change is driven by the true loss of eS25 expression, rather than other artefacts from CRISPR/Cas9 (58, 70). The failure of homology-directed repair to rescue phenotypes or gene expression markers suggests that restoring expression in the most native way possible still cannot revert cells to the wild-type state (**Figure 8**). We expect that loss of RPS25 could affect cellular differentiation in other contexts, but in some cases is lethal given that RPS25 is a common essential gene (depmap.org). Phenotypes associated with reduced levels of but not complete loss of eS25, such as resistance to ricin toxicity in K562 cells (40), may arise by a similar mechanism, especially since this cell line can differentiate in culture and genetic rescue has not been tested (71).

Prior literature has led to a near synonymous association of eS25 with IRES-mediated and specialized translation events (20, 34), and it is often assumed that eS25 has a direct effect on translation when its loss is associated with a phenotype. We postulate that these effects, with the exception of CrPV IGR IRES-mediated translation, are indirectly related to eS25’s true function. We found that the requirement of eS25 for ribosome recruitment to the CrPV IGR IRES is not generalizable to other IRES types, and neither eS25 nor RACK1 are required for HCV IRES-mediated translation (**Figures 1** and **Figure 3**). Given that the CrPV IGR IRES represents a mechanistically unique IRES, which mimics an elongation rather than initiation state and utilizes no initiation factors (72), dependence on eS25 should not be extrapolated to other translation mechanisms. The accepted role of other common essential RPs (*e.g.*, uL1/RPL10A and eL38/RPL38 (73, 74)) in directly controlling IRES-mediated translation might be revisited in this light given that requirements have not been shown *in vitro* and rescue experiments have not been tested.

We propose that the current focus on specialized translation has overlooked other critical roles for eS25 in cell biology. While it has been concluded that natural eS25 ribosome deficiency is selected by the cell to preferentially translate certain messages, we speculate that deficiency could instead be a consequence of aberrant translation events and quality control. The elevated eL22L1 and eS6-P we observed in the RPS25 knockout suggests that the cellular ribosome pool was under a stress relating to its biogenesis and turnover. Intriguingly, eS25 was recently demonstrated to interact with the Not5 subunit of CCR4-Not in budding yeast (75), and preliminary analysis of our HEK293T mutant supports this connection (**Figure S21**). Given that CCR4-Not participates in roles across gene expression and ribosome biogenesis (76, 77), we expect that perturbing such a relationship could incur broad cellular dysregulation like the ones we observe. Our RNA-seq and quantitative imaging analyses drew particular attention to nucleolar dysregulation of the RPS25 knockout (**Figure 6** and **Figure S18**). Though RPS25 has been linked to a similar stress of other orphan RPs through MDM2-p53 (78), we observed WT levels of p53 in the RPS25 knockout and no indication of p53 target gene transcriptional dysregulation by RNA-seq (**Figure 5E**). Given the specific outcomes of RPS25 loss in yeast that appear independent of ribosome levels (29), we presume that p53 dysregulation is not the major consequence of RPS25 loss and other yet to be defined mechanisms are at play. Notably, studies have found eS25 in early pre-ribosomal particles within the nucleolus, despite the finding that its depletion does not alter pre-18S rRNA maturation or ribosome levels like other “initiation RPSs” (12, 13, 79). Taken together, these observations highlight unappreciated roles of eS25 in ribosome biogenesis and quality control and motivate deeper investigation.

Finally, since it has been speculated that ribosome heterogeneity may be therapeutically targeted (22, 80), our findings provide an important word of caution. Though mutations in RPS25 are not linked to any ribosomopathies, the differentiation of the RPS25 knockout may be a general phenomenon associated with genetic lesions in ribosomal components. Indeed, our unbiased genetic screen for regulators of eS6 phosphorylation revealed that RPS25 loss is not alone amongst RPs in driving indirect effects, some of which are tied to disease and specialized translation mechanisms (**Figure 5**). If RP mutations indirectly drive cellular transformation like that for RPS25, rather than directly interfering with translation, gene therapy-mediated repair in somatic cells might ultimately fail. The therapeutic reduction of eS25 and other RPs may also cause disease if it promotes inflammation or nucleolar dysregulation (67, 74). Our study collectively uncovers complex outcomes that follow ribosomal protein loss in a human cell and shows that even in the most suggestive case, genetic customization of the ribosome does not imply translational causality.

## MATERIAL AND METHODS

### Cell growth and lentiviral transductions

HAP1 cells were grown at 37°C with 5% CO_2_ in Iscove’s Modified Dulbecco’s medium (IMDM) supplemented with 10% (v/v) heat-inactivated fetal bovine serum, 2 mM L-glutamine, and 1X penicillin/streptomycin. K562 cells were grown at 37°C with 5% CO_2_ in RPMI media with 10% (v/v) fetal bovine serum, 2 mM L-glutamine, and 1X penicillin/streptomycin. HEK293FT cells were grown at 37°C with 5% CO_2_ in DMEM media with 10% (v/v) fetal bovine serum, 2 mM L-glutamine, and 1X penicillin/streptomycin. Wild-type, RPS25 KO (CRISPR/Cas9 KO1 clone A8-15 (KO1), and gene trap insertion clone 45-1 (KO2)), RACK1 KO (CRISPR/Cas9 clones E3-A5 (KO1) and KO2 E3-A6 (KO2)), and eIF3H KO HAP1 cell lines were used in this study (9, 44, 61, 81). Cells lines all tested negative for mycoplasma with the MycoAlter PLUS Mycoplasma Detection Kit (Lonza cat.#LT07-703). Lentivirus packaging was conducted at the Gene Vector and Virus Core of Stanford University or performed in-house using lentiviral constructs by co-transfection with ΔVPR, VSV-G, and pAdVAntage packaging plasmids into HEK293FT cells using FuGENE HD (Promega) (82). Cells were transduced and selected as described (81) using constructs in the pLenti CMV PURO vectors expressing RPS25-HA, RPS25-ybbR, or RACK1-FLAG. The K562 cell lines bearing RPS25-targeting and control shRNA were gifts from G Hess and M Bassik.

### Ribosome purification and polysome profiling

Purification of human 40S and 60S ribosomal subunits and polysome profiling was performed as described (10). Purification of “crude 80S” ribosomes for mass spectrometry was performed by halting purification of ribosomal subunits after resuspending the ribosome pellet from the first sucrose cushion and prior to splitting subunits. For purification of crude 80S ribosomes from the K562 cells, the method was identical except that the cells were pelleted from suspension culture prior to lysis rather than scraping from culture dishes. For purification of monosomes (“mono”) and polysomes (“poly”) from polysome profiles, fractions corresponding to each were pooled from two profiles per cell line. These pooled fractions were then pelleted with a low-salt sucrose cushion (100 mM KOAc, 5 mM MgOAc_2_, 30 mM HEPES-KOH pH 7.5, 1 mM DTT) by centrifuging at 63,000 x *g* for 18 hrs at 4°C using a Type 80 Ti rotor (Beckman Coulter). Pellets were resuspended and these samples were analysed by mass spectrometry.

### Membrane purification

Purification of cellular membranes (sometimes called “ER membranes” or “ER”) was performed roughly as described (68). Briefly, HAP1 cell lines were grown with regular passages and seeded in single 15-cm dishes in triplicate for each cell line 48 hrs of growth before harvesting at ∼80 confluency. To normalize growth conditions prior to harvest, media was removed 6 hrs prior to harvest and replenished with fresh media. Cells were harvested by aspirating media and washing cells on plate 2X with 10 mL pre-chilled PBS. Cells were scraped in residual PBS, pelleted, and then resuspended in 1 mL permeabilization buffer (110 mM KOAc, 2.5 mM MgOAc2, 25 mM HEPES-KOH pH 7.5, 1 mM EGTA, 0.015% digitonin, 1 mM DTT) per 15-cm plate. Permeabilized cells were incubated for 5 min then centrifuge for 10 min at 1,000 xg. The supernatant was removed and saved “cytosol fraction”, and the pellet was resuspended with 5 mL wash buffer (110 mM KOAc, 2.5 mM MgOAc2, 25 mM HEPES-KOH pH 7.5, 1 mM EGTA, 0.004% digitonin, 1 mM DTT) and re-pelleted at 1000 xg for 10 min. The wash was removed and the cytosol-vacated pellet was mixed with 250 uL lysis buffer (110 mM KOAc, 2.5 mM MgOAc2, 25 mM HEPES-KOH pH 7.5, 1% NP-40, 0.5% sodium deoxycholate, 1 mM DTT) and incubated on ice for 5 min. Nuclei were pelleted at 7,500 xg for 10 min, and the supernatant (“membrane fraction”) was saved. Membrane samples were subsequently analysed by mass spectrometry.

### Gel electrophoresis and immunoblotting

Native acrylamide/agarose composite gels were cast and run as described (81). Complexes were formed with the indicated amounts of ribosomal subunits and labelled RNAs in a buffer containing 30 mM HEPES-KOH (pH 7.4), 100 mM KOAc_2_, and 2 mM MgOAc_2_, unless otherwise indicated. The HCV IRES RNA was fluorescently labelled at the 3’ end as described (10), and the CrPV IGR IRES A1F construct used previously (83) was transcribed and 3’ end labelled in a similar manner.

Except when otherwise stated, immunoblotting was performed as described (81). When blotting purified 40S ribosomal subunits, an equal concentration of 40S (by A_260_) was loaded into separate wells of the SDS-PAGE gel (10 pmol). When blotting either crude 80S ribosomes, post-nuclear cell lysate, or whole cell lysate, respective protein concentrations were determined for each sample with a Bradford assay (BioRad) such that protein concentrations could be normalized. When blotting whole cell lysates, lysate was first treated with DNAse I (NEB) to mitigate genomic DNA viscosity when loading into SDS-PAGE gels. When re-probing blots, HRP-conjugated secondary antibodies were either inactivated by incubation with 0.02% sodium azide in 5% skim milk/TBST or stripped with Restore Stripping Buffer (ThermoFisher cat.#21059). Lysate blots were re-probed with anti-p84 and/or anti-GAPDH-HRP as loading controls, while ribosome blots were blotted with antibodies against other ribosomal proteins as appropriate. Most antibodies were used in a block of 5% skim milk in TBST, except the anti-eL22L1 antibodies which were used in a block of 3% BSA in TBST. A list of antibodies used for western blots is in Supplementary Table 1.

### Dual-luciferase assays

A dicistronic vector encoding a 5’ cap-driven Renilla luciferase and downstream HCV IRES driven firefly luciferase was used for dual-luciferase assays (Fig. 2). The plasmid was transfected into HAP1 cell lines as described (81) and then cells were lysed and assayed under various conditions and time points, as described in the main text and figures. Under similar conditions, cell lines were assayed with a Propidium Iodide (PI) FACS-based assay (Abcam cat.#ab139148) to assess the ploidy of each cell line under the conditions indicated. The resulting PI assay data was analysed in FlowJo to isolate singlets and plot histograms of PI intensity using the 488nm laser with the 615/25 filter and B615 detector. All statistical analyses were performed in GraphPad Prism.

### Mass spectrometry

Purified ribosome subunits were isolated and quantified as described above. 10 μg of protein was used as input material for each digestion. Samples were brought up to 95 μL in 50 mM Ammonium Bicarbonate (AmBic) and reduced with 5 mM DTT for 20 min at 60°C. Samples were cooled to room temperature and then alkylation was achieved by adding 30 mM iodoacetamide for 30 min at 25°C in the dark. To digest peptides, 400 ng of sequencing grade trypsin (Promega) was added for 16 hrs at 37°C. Samples were subsequently acidified by adding formic acid to a final concentration of 2.5% and incubating at 37°C for 45 min. Finally, samples were desalted using HyperSep Filter Plates with a 5-7 μL bed volume (Thermo Fisher Scientific) following the manufacturer’s instructions. Samples were eluted three times in 100 μL 80% ACN in 0.2% formic acid, dried on a SpeedVac, and resuspended in 10 μL 0.2% formic acid for mass spectrometry analysis.

Cell membrane fractions from HAP1 cells were purified and quantified as described above. To prepare peptide samples, 10 μg of protein was used as input on S-trap Micro Column (Protifi) as per the manufactures protocol. To digest peptides on-column, 750 ng of sequencing grade trypsin (Promega) was added for 1 hr at 48°C. Digested peptides were eluted sequentially with 40 μL of 0.2% formic acid and then 50% ACN in water. Samples were dried on a SpeedVac and resuspended in 10 μL 0.2% formic acid for mass spectrometry analysis.

Samples were analysed by online nanoflow LC-MS/MS using an Orbitrap Fusion Tribrid mass spectrometer (Thermo Fisher) coupled to a Dionex Ultimate 3000 HPLC (ThermoFisher). A portion of the sample was loaded via autosampler isocratically onto a C18 nano pre-column using 0.1% formic acid in water (“Solvent A”). For pre-concentration and desalting, the column was washed with 2% ACN and 0.1% formic acid in water (“loading pump solvent”). Subsequently, the C18 nano pre-column was switched in line with the C18 nano separation column (75 µm × 250 mm EASYSpray (ThermoFisher) containing 2 µm C18 beads) for gradient elution. The column was held at 45°C using a column heater in the EASY-Spray ionization source (ThermoFisher). The samples were eluted at a constant flow rate of 0.3 µL/min using a 90 min gradient and a 140 min instrument method. The gradient profile was as follows (min:% solvent B, 2% formic acid in acetonitrile) 0:3, 3:3, 93:35, 103:42, 104:95, 109:95, 110:3, 140:3. The instrument method used an MS1 resolution of 60,000 at FWHM 400 m/z, an AGC target of 3e5, and a mass range from 300 to 1,500 m/z. Dynamic exclusion was enabled with a repeat count of 3, repeat duration of 10 s, exclusion duration of 10 s. Only charge states 2-6 were selected for fragmentation. MS2s were generated at top speed for 3 s. HCD was performed on all selected precursor masses with the following parameters: isolation window of 2 m/z, 28-30% collision energy, orbitrap (resolution of 30,000) detection, and an AGC target of 1e4 ions. Spectra were used to generate label-free quantitative (LFQ) intensities using MaxQuant and Perseus software, excluding reverse peptides and imputing missing values from a normal distribution (84, 85). Data from experiments with biological replicates were analysed by a two-tailed heteroscedastic t-test in Excel, after checking that the data was log2 normal. Mass spectrometry results were analysed and plotted in RStudio, significant overlapping changes between conditions were determined and plot using BioVenn (86), and ontology was performed using data from the Gene Ontology Project powered by Panther (87–89). Membrane MS and RNA-seq data was intersected using Cytoscape to produce Figure 7B (90). Supplementary Table 3 contains processed LFQ intensities from MS analyses.

### RT-qPCR

Cell lines, as indicated in the main text, were seeded into wells of a 96-well plate at 10,000 cells per well 24-48 hrs prior to lysis, reverse transcription, and amplification with the Cell-to-C_T_ kit (ThermoFisher cat#4402955). Real-time PCR was performed with the SYBR Green PCR master mix (ThermoFisher cat#4309155) using a CFX Connect Real-Time System (BioRad). In experiments with tunicamycin-treated cells, tunicamycin or DMSO was added to wells 24 hrs post-seeding, and then cells were lysed and assayed after 24 hrs in the presence of drug. Primers used for RT-qPCR are listed in Supplementary Table 2 and data was plot in GraphPad Prism.

### Viral infections

All viral infections were performed under appropriate biosafety conditions using viruses titred by standard plaque assays. The DENV-luc experiments used virus produced from BHK-21 cells, by transfection of an RNA encoding the HAP1-adapted dengue serotype 2 virus (DENV-2) with a Renilla luciferase ORF embedded at the 5’ end of the DENV ORF (45). The Coxsackie B3 luciferase (CV-B3-luc, Nancy strain) experiment used virus produced by transfection of the infectious clone pRLuc-53CB3/T7 into RD cells (91). HAP1-adapted DENV-2 virus (clone 16681) was propagated in C6/36 cells or HAP1 cells, Chikungunya virus (CHIKV, 181/25 vaccine strain) was propagated in BHK-21 cells, and Zika virus (ZIKV, PRVABC59 (Human/2015/Puerto Rico, NR-50240)) was propagated in C6/36 cells.

For most DENV-luc infections, cells were seeded into 96-well plates at 10,000 live cells/well, infected 24 hrs after seeding, and then lysed and assayed 24, 48, and/or 72 hrs post-infection. Except when otherwise stated, cells were infected with a low MOI of 0.018. When indicated, cells were treated with the replication inhibitor MK0608 or translation inhibitor cycloheximide (CHX) at 50 μM and 100 μM, respectively. For adherent cells, media was directly aspirated from wells at indicated time points, followed by the addition of 100 uL Renilla Luciferase lysis buffer (Promega cat.#E2810), and appropriate incubation. Renilla luciferase units (RLU) were measured either by the addition of 20 uL lysate to 100 uL Renilla luciferase assay reagent in an eppendorf tube and measuring luminescence using a GloMax 20/20 Single Tube Luminometer with 5 sec integrations, or by sequential addition of 50 uL Renilla luciferase assay reagent into wells containing 20 uL lysate and measuring luminescence via a 5 sec integration using a Veritas microplate luminometer in white 96-well plates (Corning cat.#3789A).

For crystal violet staining of cells upon virus infections, cells were seeded at 10,000 live cells per well in a 96-well plate and infected with viruses at the indicated MOIs. At 3 days post-infection, cells were fixed with 4% paraformaldehyde in PBS and viable cells were visualized by crystal violet staining. To titre the production of DENV-2 infectious particles by plaque assay, HAP1 cell lines were seeded into 6-well plates in triplicate at 250,000 lives cells per well. At 24 hrs post-seeding, cells were infected with DENV-2 at MOI=0.1 PFU/cell, and the infection was allowed to proceed for 48 hrs. At the end of the experiment, media from infected cells was collected, and cells were washed with PBS and then lysed on plate with RIPA buffer. The DENV-2 secreted into supernatants harvested from infected cells were subsequently tittered by plaque assays using Huh7.5.1 cells, while the cell lysate was used for immunoblotting with antibodies against DENV proteins (52). All statistical analyses were performed in GraphPad Prism.

### Proliferation assays

Proliferation assays were performed roughly as described (81), and modified to include select drug treatments. Briefly, HAP1 cell lines from routine passages were dissociated from culture plates with trypsin and counted for live cells by trypan blue. Cells were then seeded into 96-well plates at 10,000 live cells/well with 3-10 replicates per cell line. 24 hrs after seeding cells, drugs or DMSO was added to respective wells via 2X stocks in media, and then cells were assayed over 4 days of growth using the MTT reagent (ThermoFisher). Since tunicamycin does not have a discrete molecular weight, we approximated it as 840 g/mol to facilitate ease in plotting molar concentrations in Figure 4 (*e.g.*, 0.25 μg/mL ≈ 300 nM). Absorbance values at 570 nm were determined using a Synergy Neo2 instrument (BioTek). All statistical analyses were performed in GraphPad Prism.

### Immunofluorescence imaging and analysis

Wild-type, RPS25 KO1, and RPS25 KO2 HAP1 cell line were seeded into a glass bottom 96-well plate (Corning cat.#4580) at 10,000 live cells per well and incubate under standard conditions. 24 hrs after seeding, a portion of cells were infected with HAP1-adapted DENV-2 at MOI=2. The infection was allowed to proceed for 24 hrs, at which point media was aspirated from wells and cells were fixed with 4% PFA in PBS for 15 min at room temperature. Following three washes in PBS, cells were permeabilized with 0.1% triton X-100 in PBS for 15 min, washed again 3X with PBS, and then blocked overnight at 4°C with 5% BSA in PBS. Cells were either immediately stained with antibodies or kept in PBS containing 0.02% sodium azide at 4°C until use. Cells were stained with antibodies as indicated in the main text and figures, by first incubating in primary antibodies overnight in PBS with 2% BSA at 4°C. Cells were then washed 3X with PBS, incubated in PBS with 2% BSA for 2 hrs at 4°C, and washed 3X with PBS prior to imaging. Concurrently with secondary antibody staining, cells were stained with Hoescht 33342 (Invitrogen cat.#H3570) at 0.5 μg/mL and Phalloidin 660 (Invitrogen cat.#A22285) from a methanol stock for a final concentration of 80 nM. A list of antibodies used for immunofluorescence staining is in Supplementary Table 1.

Epi-fluorescent imaging was performed with an ImageXpress Micro XLS Widefield High-Content Analysis System (Molecular Devices) using a 20x Plan Apo objective with 2x camera binning. Confocal imaging was performed with a Nikon A1R HD25 microscope and 60X oil objective using 405, 488, 561, and 640 nm lasers as appropriate. Multiple Z-stacks were acquired for each condition by confocal, and for comparing images across conditions Z planes that transected similar regions of cells were chosen. For comparative image analysis, only images acquired with the same laser settings were compared and the antibody channel intensity was maintained constant between conditions when preparing images for figures.

For quantitative image analysis, cellular nuclei were first segmented based on Hoechst staining. Nuclear images were converted to a mask using the minimum error thresholding method. Touching cells were then split using two steps applied in serial. The first step used a marker control watershed approach where the markers were derived from regional maximum values of the nuclear image. In the second step, the watershed image was used to further split cells in contact with their nearest neighbor(s). A custom segmentation algorithm was implemented to detect and bridge concave inflections in the perimeter of each object to separate any remaining touching cells. Resulting objects that were too small, too large, or oddly shaped were not included in further analysis. For signal measurement from each signal channel, a large radius top-hat filter was first applied to subtract the background signal. The nuclear mask image was then used to mark regions of interest to calculate the signal intensity within each cell nucleus. Nuclear immunofluorescence signals were calculated as mean nuclear intensity of the pixels in each cell nucleus. Image analysis was performed using custom scripts written in Matlab (available at Github).

### Transmission electron microscopy

Wild-type, RPS25 KO1, and RPS25 KO2 HAP1 cell lines were seeded into a 6-well plate at 500,000 live cells per well. Prior to seeding, a 10×5mm Aclar slab (Ted Pella cat.#10501-10) was deposited in wells to which cells adhered after standard incubation. 24 hrs after seeding, a portion of cells were infected at MOI=2 and the cells were returned to incubation. Following 24 hrs of infection, infected and uninfected cells were fixed by rapidly transferring Aclar slabs into Karnovsky’s fixative (2% Glutaraldehyde (EMS cat.#16000) and 4% PFA (EMS cat.#15700) in 0.1M Sodium Cacodylate (EMS cat.#12300) pH 7.4) for 1 hr, chilled and delivered to Stanford’s CSIF on ice. Slabs were then post-fixed in cold 1% Osmium tetroxide (EMS cat.#19100) in water and allowed to warm for 2 hrs in a hood, washed 3X with ultra-filtered water, then all together stained for 2 hrs in 1% Uranyl Acetate at room temperature. Samples were then dehydrated with a series of ethanol washes for 10 min each at room temperature beginning at 50%, 70%, 95%, changed to 100% 2X, then Propylene Oxide (PO) for 10 min. Samples were then infiltrated with EMbed-812 resin (EMS cat.#14120) mixed 1:1, and 2:1 with PO for 2 hrs each. The samples were then placed into EMbed-812 for 2 hrs, opened and then placed into flat molds with labels and fresh resin and placed in a 65°C oven overnight.

Sections were taken around 90nm, picked up on formvar/Carbon coated slot Cu grids, stained for 40 sec in 3.5% Uranyl acetate in 50% acetone followed by staining in 0.2% Lead Citrate for 6 min. Grids were images in a JEOL JEM-1400 120kV microscope and photos were acquired using a Gatan Orius 2k X 2k digital camera.

### Genome-wide CRISPR-Cas9 screening

Immunoflow conditions were optimized using wild-type and RPS25 KO1 HAP1 cells to verify the sensitivity of an anti-eS6-P antibody (Cell Signalling #2211). Briefly, wild-type and knockout cells grown in parallel 15-cm dishes were harvested by trypsinization and counted by trypan blue staining. To normalize growth conditions prior to harvest, media was removed 6 hrs prior to harvest and replenished with fresh media. An equal number of cells (10 million) was resuspended with Cytofix/Cytoperm buffer (BD Biosciences cat.#554655) for 30 min at room temperature. Fixed cells were washed three times with 1X Perm/Wash solution (BD Biosciences cat.#51-2091KZ) and then incubated with diluted primary antibody in 1X Perm/Wash solution overnight at 4°C. After three washes in 1X Perm/Wash solution, cells were resuspended with diluted secondary antibody for one hour at room temperature in the dark. Cells were then washed three times with 1X Perm/Wash solution, passed through a 70 μm strainer (Falcon cat.#352350), and analysed by flow cytometry with the 488nm laser and 525/50 filter and B525 detector. Data was analysed with FlowJo (version 10.6.2) to isolate singlets and determine the spread of signal between wild-type and knockout cells.

To make a mutagenized cell population for screening, the HAP1-derived eHAP cell line was transduced with the Brunello CRISPR/Cas9 library and saved in batches (92, 93). Specifically, eHap cells were stably transduced with lentiCas9-Blast and subsequently selected using Blasticidin. Next, 300 million eHap cells that constitutively express Cas9 were transduced with lentiGuide-Puro from the Brunello library at an MOI of 0.3. Cells were then selected with puromycin, expanded to 3 billion cells, and then pooled together and cryofrozen in aliquots. 100 million cells were thawed constituting over 1000X genome coverage worth of mutagenized library. These cells were expanded into twenty T175 flasks and grown to ∼80% confluence (∼1.5 billion cells total). Cells were harvested by trypsinization, counted by trypan blue staining, and 500 million cells were fixed and stained with antibody as above. Stained cells were passed through a 70 μm strainer and detected by flow cytometry using the 488nm laser as above. The high (top) and low (bottom) ∼2% of eS6-P cells were sorted into separate tubes containing 1X Perm/Wash solution using a 70 μm nozzle. Genomic DNA (gDNA) was extracted from the top (5.61 million cells) and bottom (4.87 million cells) fractions using the QIAamp DNA mini kit (QIAGEN cat.#51304). Fixed cells were decrosslinked by overnight incubation in 200 µL of PBS with added Proteinase K and 200 µL buffer AL (Qiagen) at 56°C with agitation. Two rounds of PCR were used to first amplify the guide RNA sequences from the gDNA and then to add barcodes for amplicon sequencing. The PCR products were purified from 2% agarose gels via the QIAqick gel extraction kit (QIAGEN cat.#28704) and subjected to next-generation sequencing on a HiSeq instrument lane (Illumina) by Novogene. The sequencing data were analysed using the MAGeCK algorithm (version 0.5.4) and plotted in RStudio (94). A population of 5 million unsorted (unsrt) cells was analysed in parallel, and all populations were compared against the sequencing of the parental library that matched ideal library qualities (95). To control for guide RNA enrichment due to outgrowth of the parental library, genes enriched in the unsrt library (P<0.001, FC>2.5) were excluded from plots and analyses of enriched genes from eS6-P screen. Gene ontology analysis was performed using data from the Gene Ontology Project powered by Panther (87–89). Processed MAGeCK analysis results are reported in Supplementary Table 4 and antibodies are listed in Supplementary Table 1.

### RNA-seq

HAP1 cell lines were seeded into the wells of 6-well plates with equal live cell counts (250,000 cells/well), grown under standard conditions, and harvested by trypsinization, pelleting and washing cells with PBS 48 hrs post-seeding. Prior to harvest, well were replenished with fresh media as above. RNA was purified from cells pellets using the PureLink RNA mini kit (ThermoFisher cat.#12183025). Prior to library preparation, RNA concentration and sample quality by RNA integrity number (RIN) were checked using Tapestation RNA ScreenTape reagents (Agilent cat.# 5067-5576). cDNA libraries were prepared with the SureSelect Strand-Specific RNA Library Prep Kit (Agilent cat.#G9691B) on an Agilent Bravo Automated Liquid Handling Platform accordingly to the protocol (Version E0, March 2017, G9691-90030). Library concentration and integrity were checked using Tapestation D1000 ScreenTape reagents (Agilent cat.#5067-5582) and the Qubit dsDNA BR Assay Kit (Invitrogen cat.#Q32850). Sequencing was performed on an Illumina HiSeq 4000 with 2×101 base pair reads and Illumina Single Index. Reads were aligned to the hg38 reference genome using STAR v2.5.3a (96) and differential expression between samples was computed using R v3.4.0 and the DESeq2 package (97). In the rare case that fold change P-values were too small for R to calculate and estimated as 0, these were imputed at 10^−300^ to facilitate logarithmic conversion. Most graphics were generated in RStudio, Venn diagrams were prepared using BioVenn (Hulsen et al., 2008) or Venny (Oliveros, 2007), and gene ontology analysis was performed as above. Hierarchical clustering was performed by centroid linkage in Cluster 3.0 (98) and heatmaps were visualized in and exported from Java TreeView (99). We expect that RPS25 mRNA was at reduced levels in the knockout cells due to transcript degradation by nonsense-mediated decay or premature transcriptional termination, and that the RPS25-HA transcript was not detected since our library preparation relied on polyA enrichment and the lentiviral expression instead has a WPRE. Supplementary Table 5 contains processed RNA-seq data with fold-change and P-values for each condition versus WT.

### RAN translation experiments

The HA-tagged polyGA dipeptide was expressed from a C9orf72 66-repeat (C9-66R) expression construct, as previously described (22). Briefly, WT, RPS25 KO1, and RPS25 KO1 expressing RPS25-ybbR (eS25-ybbR AB) HAP1 cells were transfected with appropriate constructs using Lipofectamine 3000 and assayed in parallel 72 hrs post-transfection. Transfected cells were lysed with radioimmunoprecipitation assay buffer containing 1X HALT protease (Pierce) and post-nuclear lysate was prepared by centrifugation at 10,000xg for 10 min at 4°C. The lysate was quantified with a BCA assay (Pierce) and equal amounts (20-25 μg) were analysed by immunoblotting as described (22). In this experiment, the blot was cut into three appropriate sections and blotted for each of three antibodies. Three biological replicates of the RPS25-ybbR rescue experiment were performed (two shown) each with similar results.

### CRISPR/Cas9 mutagenesis and homology-directed genome repair

To generate a RPS25 mutant in HEK293T cell lines, we transfected cells with a vector encoding a guide targeting RPS25 exon 2. Guide strand oligos were cloned into the PX458 vector (100) using digestion-ligation with the BpiI enzyme. 48 hrs post-transfection, GFP-positive clones were sorted into 50% conditioned media and screened for eS25 by immunoblotting. Potential clones were subsequently validated by sequencing amplicons from genomic DNA.

To repair the six nucleotide deletion at the RPS25 locus in a previously-reported HAP1 knockout clone (9), we designed CRISPR/Cas9 guide strands targeting sequences upstream and downstream of the deletion, as well as homology templates with the parental sequence (**Figure S19**). The repair templates were ordered as gene blocks (IDT) and PCR amplified with Phusion polymerase (NEB cat.#M0503S). The RPS25 KO1 cells were seeded into the well of a 6-well plate at 250,000 cells/well and 24 hrs post-seeding the PX458 plasmids and respective homology templates were co-transfected into the RPS25 KO cells using Lipofectamine 3000 (ThermoFisher cat.#L3000075). Transfection was performed following the manufacturer’s protocol using 2 μg of each plasmid and template. Cells were dissociated from plates 48 hrs post-transfection, passed through a 70 μm strainer, and single GFP-positive cells were sorted into 96-well plates containing 50% conditioned media. Clones were identified ∼2 weeks after sorting and screened by genomic DNA extraction with Quickextract (Lucigen cat.#QE09050) and parental allele-specific PCR probes and GoTaq Green Master Mix (Promega cat.#M712). Clones that tested positive by PCR were further validated by western blotting for eS25, followed by amplifying gDNA fragments with Platinum PCR Supermix HiFi (ThermoFisher cat.#12532016) and Sanger sequencing. Validated clones were characterized by dengue virus infections, as described above. All oligos used for guide strand cloning, repair template production, and gDNA screening are listed in Supplementary Table 2.

## AVAILABILITY

Scripts for RNA-seq and quantitative imaging analysis are available on Github (https://github.com/emc2cube/Bioinformatics/ and https://github.com/algejo/HAP1_nuclei_analysis).

## ACCESSION NUMBERS

The RNA-seq data has been deposited at NCBI GEO (Accession: GSE139243) and raw deep sequencing data for CRISPR-Cas9 screens is being deposited at ArrayExpress (Accession: TBD).

## ACKNOWLEDGEMENT

The authors thank Onn Brandman and Ron Kopito for discussions and comments on the manuscript; Gaelen Hess, Mike Bassik, and Nick Ingolia for sharing unpublished data and reagents; Gaby Fuchs for sharing the RPS25 expression vectors; Chris Walczak, Karim Majzoub, and Puglisi, Carette, and Bertozzi lab members for input. We thank Joe Qi at the UCSF Cytogenetics lab for karyotyping analysis; John Perrino and the Stanford CSIF for assistance with transmission electron microscopy (NCRR ARRA Award #1S10RR026780-01); Meredith Weglarz and Lisa Nichols at the Stanford FACS facility for assistance with flow cytometry; and the GSSC for high-throughput sequencing (NIH award S10OD020141). We thank the SRRC for providing computational resources and support via the Sherlock cluster.

