## Supplementart Figures for "A memory of RPS25 loss drives resistance phenotypes"

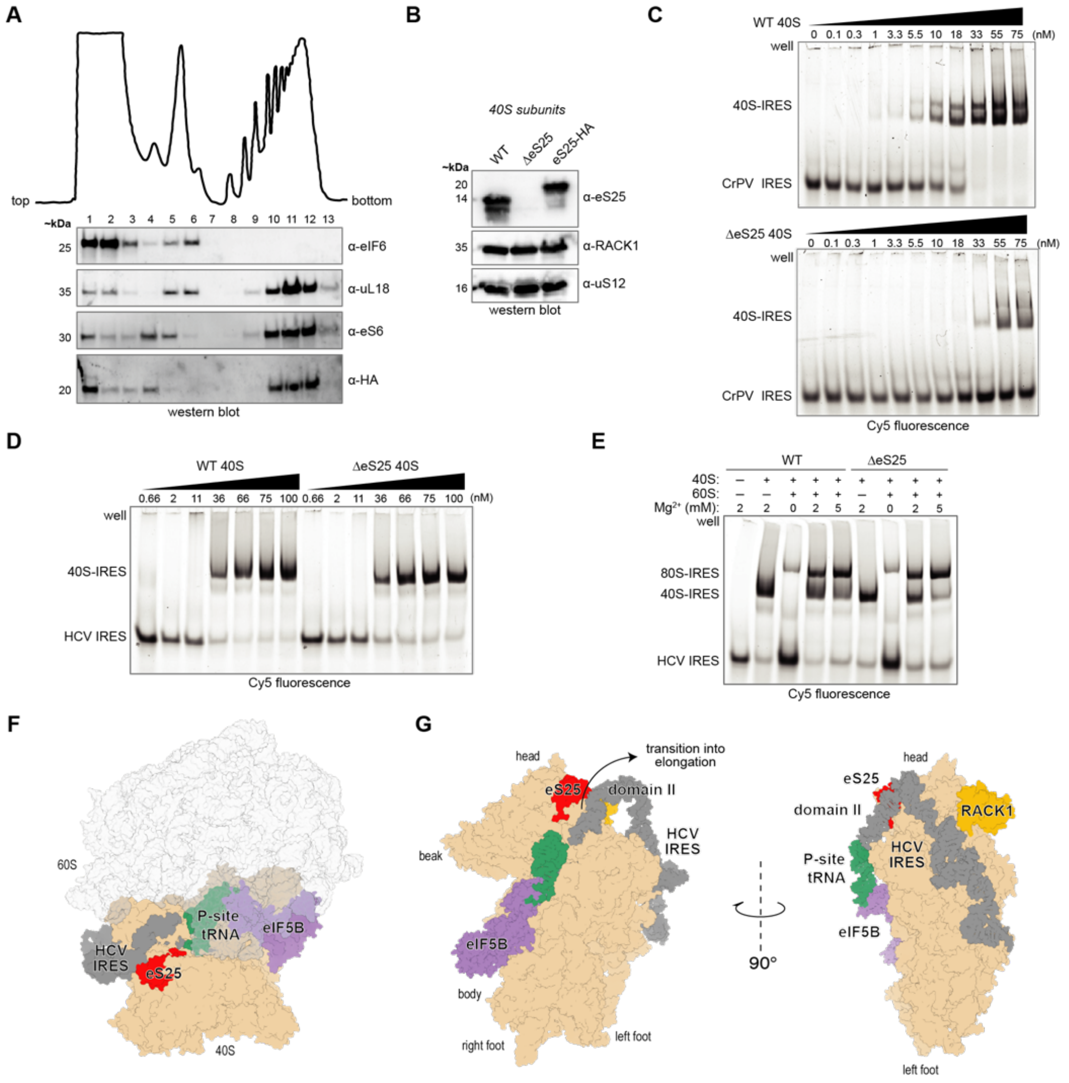

**Figure S1.** eS25 is not required for ribosome recruitment to the HCV IRES and eS25-HA functional incorporates into ribosomes. **A.** Polysome profile trace from RPS25 KO HAP1 cells that have been transduced with a lentiviral vector encoding the RPS25-HA cDNA (eS25-HA addback (AB)). Post-nuclear lysate was sedimented in 10-60% sucrose gradients, the top and bottom of the gradients are indicated, and the height of peaks is proportional to the absorbance at 254 nm. Fractions were probed for various translation proteins by immunoblotting analysis. The HA antibody used here is from Abcam. **B.** Western blot of 40S ribosomal subunits confirms successful incorporation of eS25-HA into human ribosomes. The same ribosome samples were used for gel shifts in Figure 1B. **C.** Binding of WT and ΔeS25 40S ribosomes to the CrPV IGR IRES by native gel electrophoresis. All binding reactions included 30 nM fluorescently labeled IRES RNA. **D.** Binding of WT and ΔeS25 40S ribosomes to the HCV IRES by native gel electrophoresis. Gels as in (C) but with fluorescently labeled HCV IRES RNA. **E.** Formation of 80S-HCV IRES complexes under high magnesium concentrations. Complexes were formed using 60 nM 40S with or without 120 nM 60S and resolved on an acrylamide-agarose composite gel. **F.** Structural model depicting the position of eS25 with respect to the large and small ribosomal subunits in the 80S-HCV IRES-eIF5B-Met-tRNAi-GMPPNP complex (PDB 4ujd). The small subunit is colored tan, the large subunit white, eS25 red, RACK1 is orange, the HCV IRES gray, P-site Met-tRNAi green, and eIF5B purple. **G.** Reorientation of the 80S-IRES model from (F) omitting the 60S subunit to show a similar 40S orientation as in Figure 1A. Left orientation shows the 40S subunit interface, indicating factor and tRNA positions and the rearrangement of HCV IRES domain II during the transition to elongation. The right panel shows the solvent-exposed surface of the 40S subunit (or backside) with the position of RACK1 at the head region making no direct contacts to the HCV IRES.

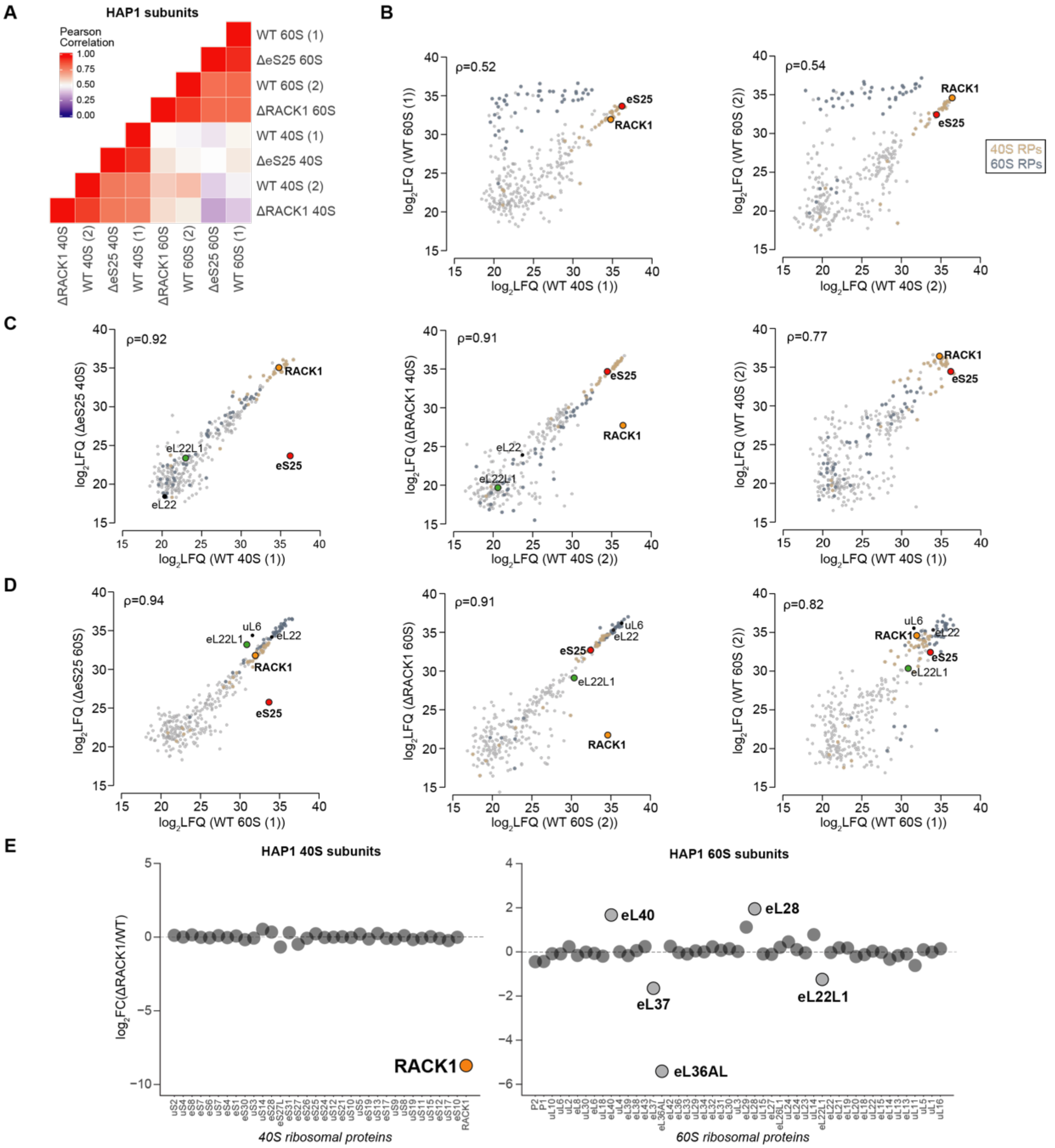


**Figure S2**. Mass spectrometry of eS25 and RACK1-deficient ribosomes. **A.** Heatmap of Pearson correlation coefficients (ρ) for $\log_2$LFQ intensities from each HAP1 ribosomal subunit samples. WT (1) was purified in parallel (paired) with the ΔeS25 sample, while WT (2) was paired to the ΔRACK1 sample. **B.** Plot of log2 LFQ intensities of 60S versus 40S subunits for WT (1) and (2) samples. **C.** Plots of log2 LFQ intensities from mutant versus WT 40S subunit samples isolated from RPS25 KO1 (ΔeS25) and RACK1 KO1 (ΔRACK1) cell lines in paired experiments. **D.** Plots of log2 LFQ intensities from mutant versus WT 60S subunit samples isolated from RPS25 KO1 (ΔeS25) and RACK1 KO1 (ΔRACK1) cell lines in paired experiments. For B-D, RPs are colored as indicated and select ribosomal and non-ribosomal proteins are annotated. **E.** Plot of the log2 fold-change in LFQ intensities between ΔRACK1 and WT ribosomal subunits. Plot is as in Figure 3A with select outlier RPs annotated.

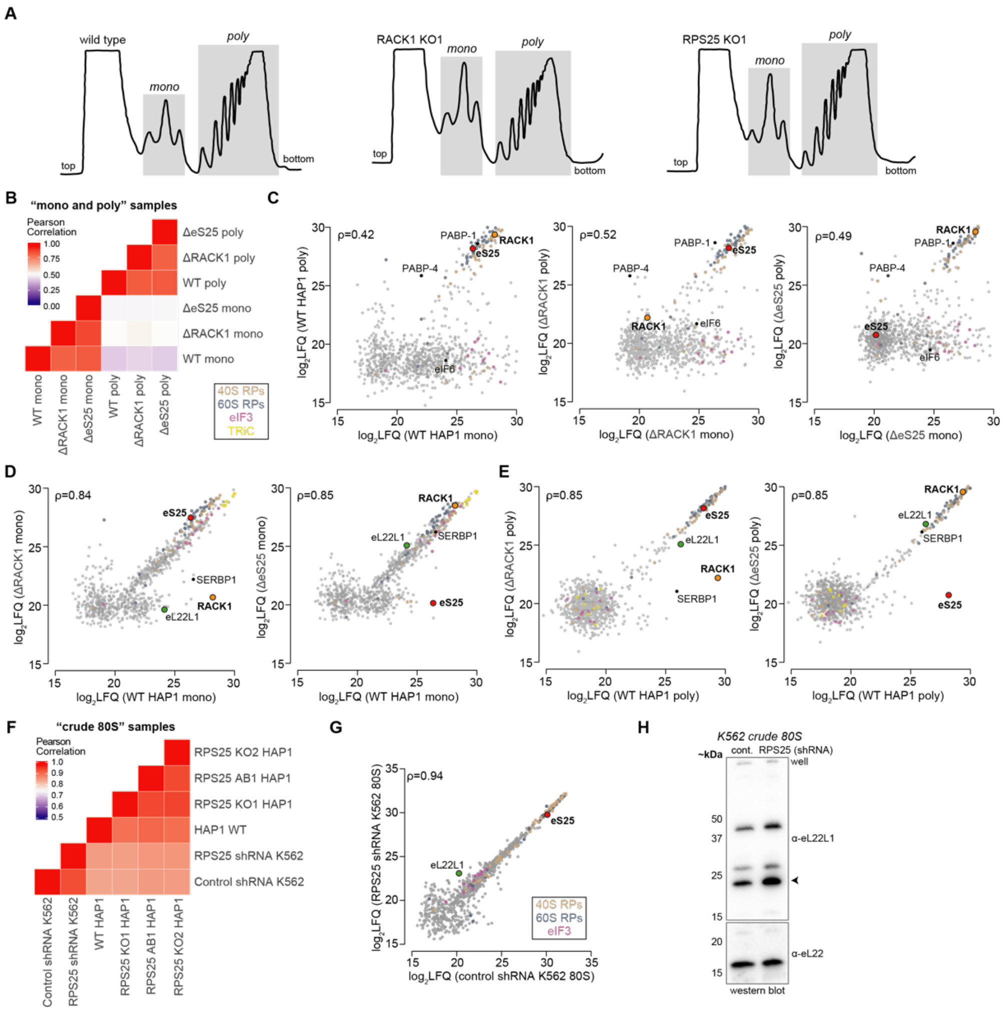


**Figure S3**. Mass spectrometry of eS25- and RACK1-deficient monosome, polysome, and crude 80S samples. **A.** Example polysome profiles of WT, RACK1 KO, and RPS25 KO HAP1 cells that were used for isolating monosomes ("mono") and polysomes ("poly") for mass spectrometry. Post-nuclear lysate was sedimented in 10-60% sucrose gradients, the height of peaks is proportional to absorbance at 254 nm, and the top and bottom of gradients are indicated. Mono and poly were isolated from fractions under respective gray highlighted regions for two polysome profiles for each cell line. **B.** Heatmap of Pearson correlation coefficients (ρ) for HAP1 mono and poly samples. **C.** Plot of log2 LFQ intensities of poly versus mono samples from WT and mutant cells. RPs and eIF3 subunits are colored as indicated and select ribosomal and non-ribosomal proteins are annotated. **D.** Plot of log2 LFQ intensities of mutant versus WT mono and poly samples. RPs, eIF3 subunits, and TRiC chaperonin complex subunits are colored as indicated. Select ribosomal and non-ribosomal proteins are annotated. **E.** Heatmap of Pearson correlation coefficients (ρ) for log$_2$LFQ intensities from each HAP1 or K562 "crude 80S" ribosome sample. **F.** Plot of log2 LFQ intensities of crude 80S ribosomal samples from K562 cells. RPs and eukaryotic initiation factor 3 (eIF3) subunits are colored as indicated and select ribosomal and non-ribosomal proteins are annotated. **H.** Western blot analysis of crude 80S subunits for eL22L1 from K562 cell line. Arrows points to the antibody-sensitive band at ~23 kDa, which is larger than expected for the protein product (~15 kDa). The identity of the higher MW bands was not further examined, but it was not present in the purified subunits (Figure 2C).

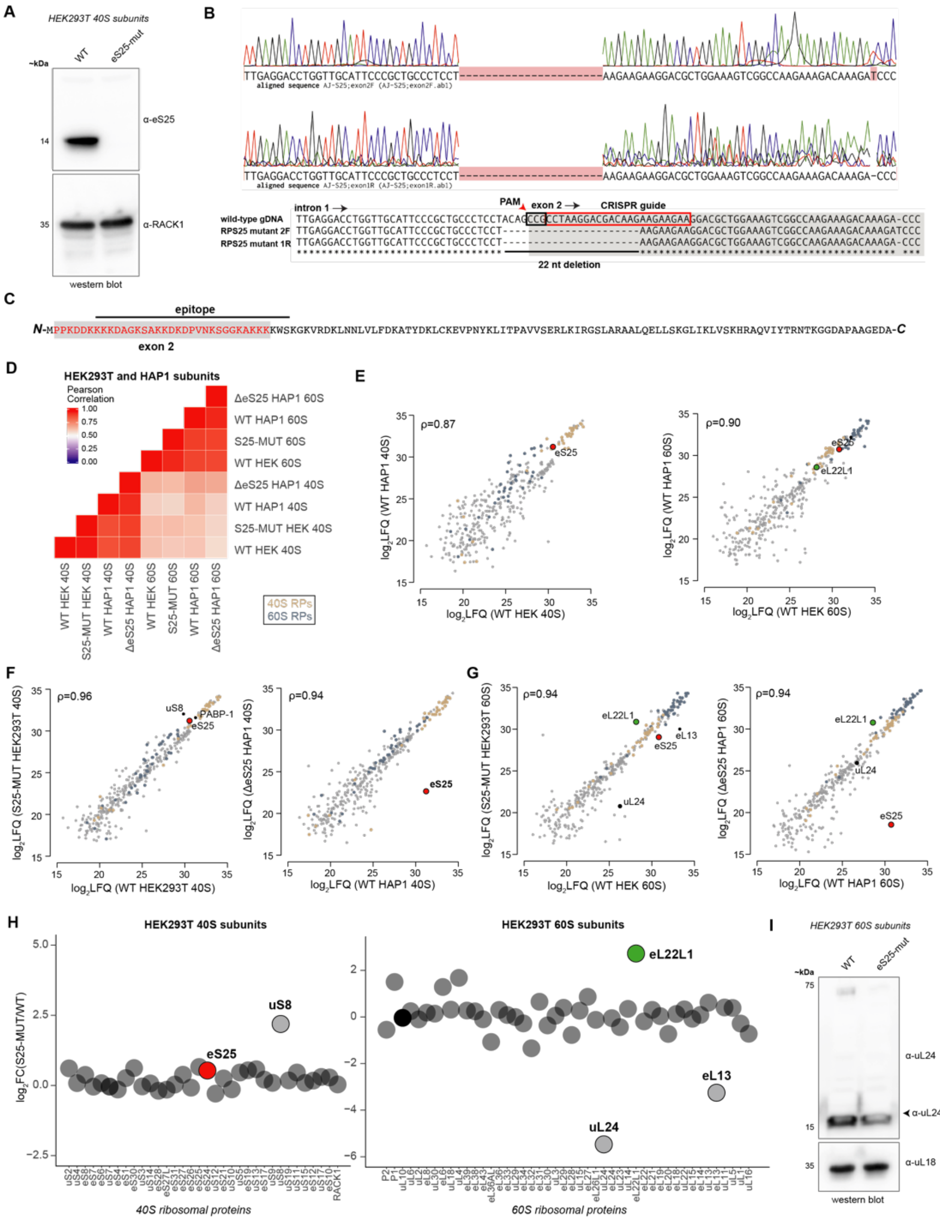

**Figure S4**. Isolation and characterization of an N-terminal RPS25 mutant (eS25-mut) from HEK293T cells by CRISPR/Cas9. **A.** Western blot analysis of WT and eS25-mut 40S ribosomal subunits demonstrates the absence of an antibody sensitive band. **B.** Genotyping of the eS25-mut cell line identified the presence of a 22-nt deletion spanning the intron 1 and exon 2 boundary. **C.** The eS25-mut may lead to loss of an antibody sensitive band via exon 2 exclusion given the N-terminal epitope. **D.** Heatmap of Pearson correlation coefficients (ρ) for $log_2$LFQ intensities from each ribosomal subunit sample. HAP1 samples (analyzed in Figure S4) were reanalyzed in parallel with the HEK293T samples. **E.** Plots of log2 LFQ intensities from WT 40S and 60S subunit samples between cell lines. **F.** Plots of log2 LFQ intensities from mutant versus WT 40S subunit samples. **G.** Plot of the log2 fold-change in LFQ intensities between mutant vs WT 60S ribosomal subunits. For E-G, RPs are colored as indicated and select ribosomal and non-ribosomal proteins are annotated. **H.** Plot of relative LFQ intensities between eS25-mut and WT HEK293T ribosomal subunits. The lack of a reduction in relative eS25 levels in the HEK293T mutant supports the occurrence of exon 2 exclusion, rather than complete loss. Plot is as in Figure 2A with select outlier RPs annotated. **I.** Western blot of 60S subunits to confirm the reduction of uL24 observed in (H), which was not further examined.

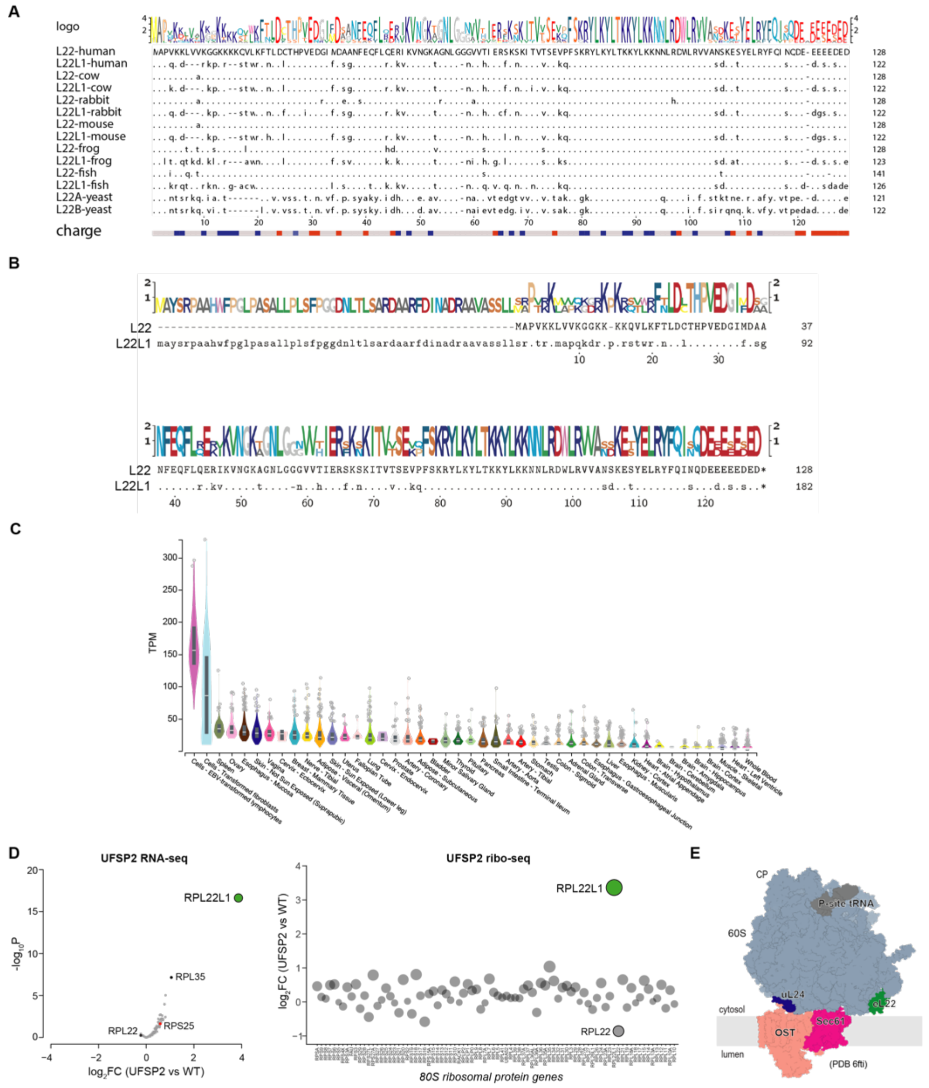


**Figure S5.** eL22L1 is distinct from its paralog and its expression correlates with cellular transformation and ribosome stress. **A.** Protein sequence alignments of eL22 and eL22L1 proteins from human (*H. sapiens*), cow (*B. taurus*), rabbit (*O. cuniculus*), mouse (*M. musculus*), frog (*X. laevis*), fish (*D. rerio*), and budding yeast (*S. cerevisiae*) retrieved from UniProt. The human eL22L1 protein sequence used here represent the short isoform without an N-terminal segment distinct from eL22, and these two proteins are 69% identical. **B.** Alignment of a long human eL22L1 isoform with eL22, based on predicted proteins from translation of ORFs predicted from UCSC Genome Browser (hg38). These two proteins are 49% identical. **C.** Expression of RPL22L1 mRNA in various tissues from the GTEx Consortium measured as transcripts per million (TPM). The highest levels of expression occur in EBV transformed lymphocytes and transformed fibroblasts. **D.** Data from a published HEK293 UFSP2 KO RNA-seq and ribo-seq dataset was used to produce a RNA-seq volcano plot (left) and ribo-seq fold-change plot (right) for RPs (1). For RNA-seq, $-log_{10}P$ represents the negative log10 of the FDR-adjusted P-value. For the ribo-seq plot, the size of the bubbles is proportional to the absolute value of the y-axis. **E.** Structural model for the position of eL22 within the ER-bound 60S-Sec61-OST complex (PDB 6fti). Gray rectangle represents the ER membrane with OST and Sec61 translocon complexes spanning. Select RPs of the 60S ribosomal subunit (eL22 and uL24—the target of ufmylation) are indicated, as well as P-site tRNA and the central protuberance (CP).

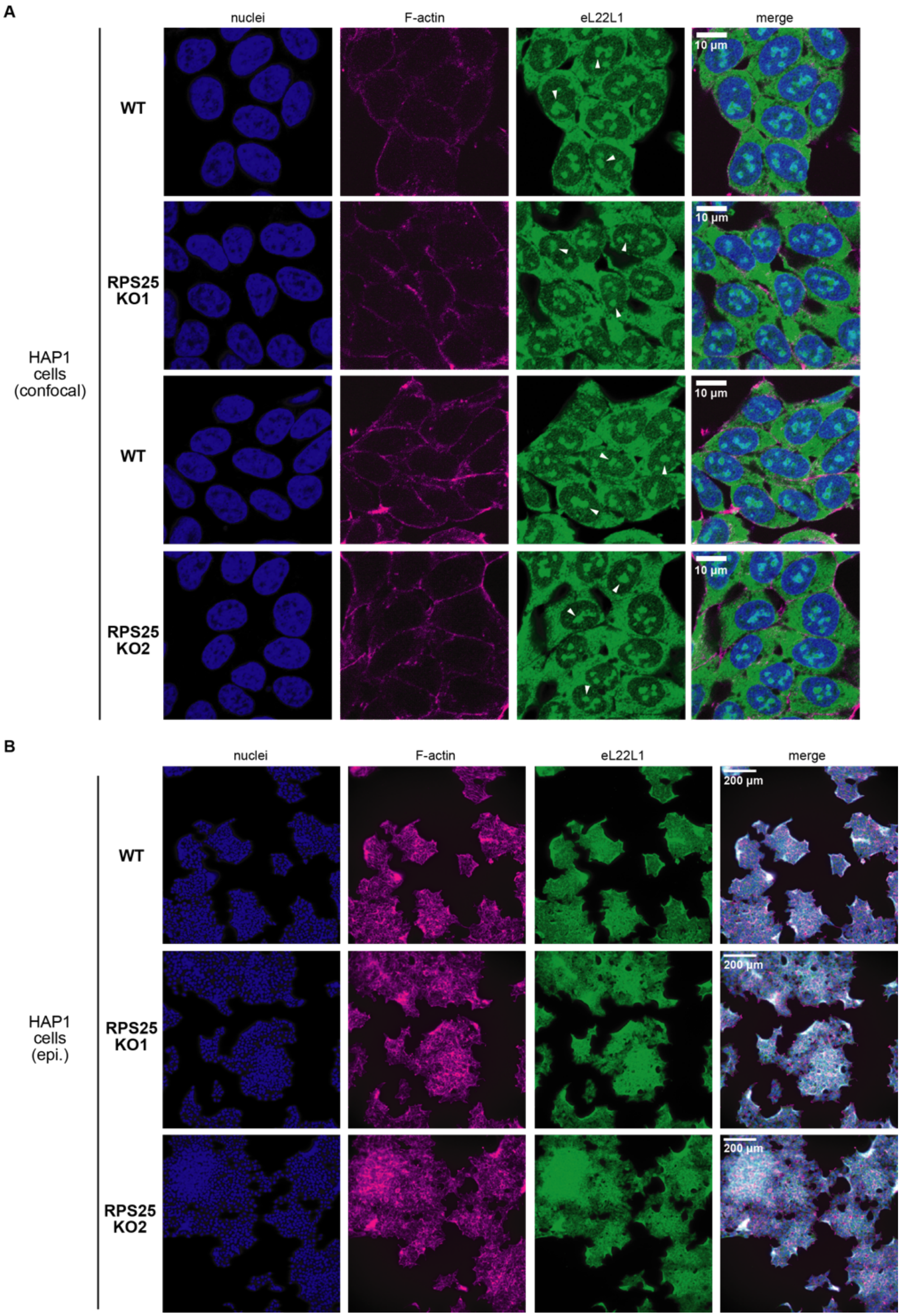


**Figure S6.** Immunofluorescent imaging of WT and RPS25 KO HAP1 cells for eL22L1. Cells were fixed and then stained with an antibody targeting eL22L1, as well as the Hoescht stain for nuclei and Phalloidin 660 for F-actin. **A.** Confocal imaging of WT and RPS25 KO2 HAP1 cells (top two panels are the full set of images from Figure 2B). White arrows point to the nucleolar staining of eL22L1 within cells. **B.** Epi-fluorescent imaging of WT and RPS25 KO cells to demonstrate widefield images of the same samples imaged by confocal microscopy.

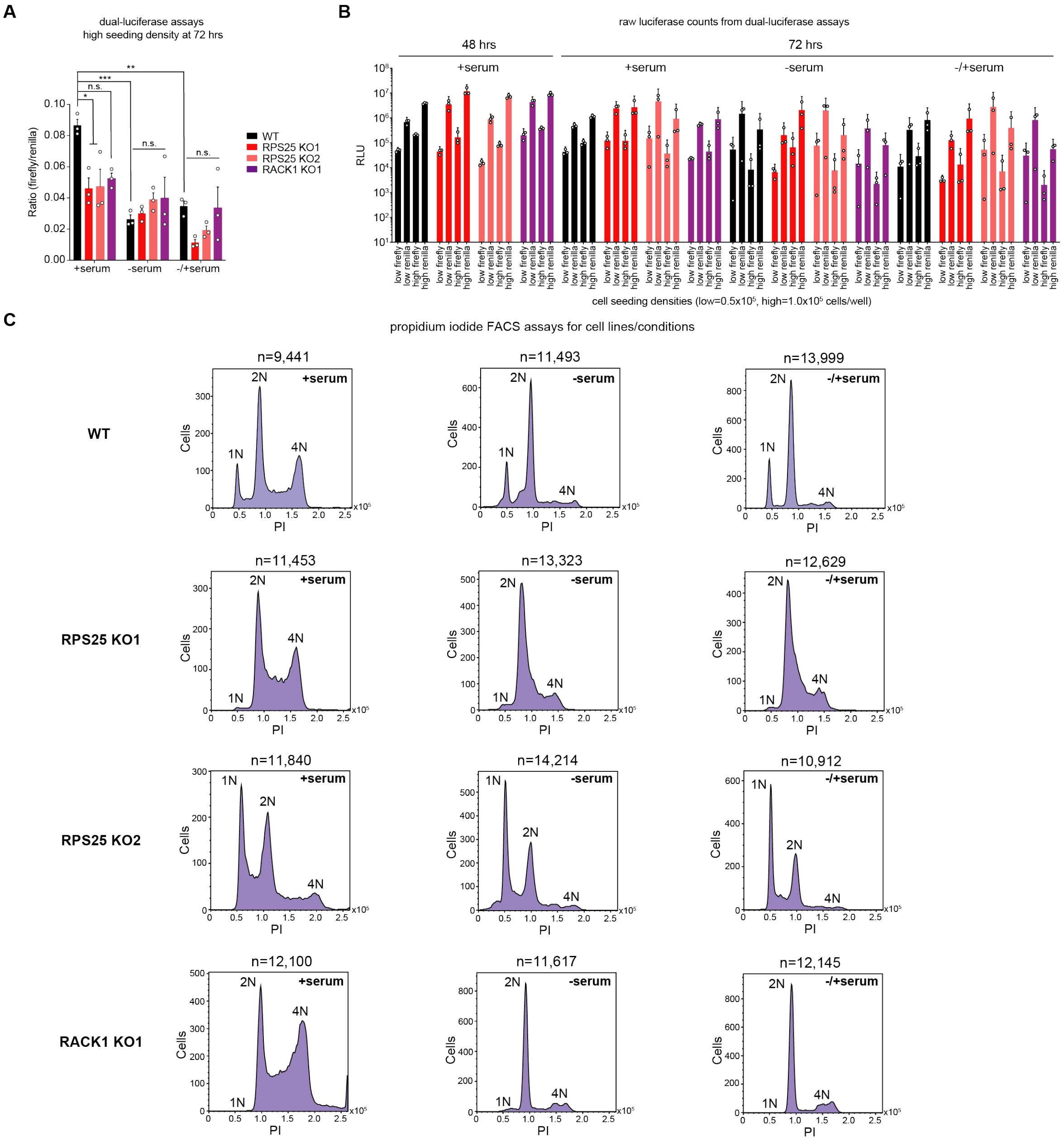


**Figure S7.** Complete results from dual-luciferase assays and validation of cell synchronization by serum starvation. **A.** Results from dual-luciferase assays of WT and KO cells at 48 hours post-transfection at the high seeding density ($1.0x10^5$ cells/well), under different perturbations of serum conditions. Error bars represent the SEM of three biological replicates and statistical comparison is the same as in Figure 2B-C. P-values: ≥ 0.05 (n.s.), 0.01-0.05 (*), 0.001-0.01 (**), and 0.0001-0.001 (***). **B.** Raw luciferase values from all experiments, subsequently used for ratio determinations in Figure 2B-C and (A). Error bars represent the 95% confidence interval for three biological replicates. **C.** Propidium iodide (PI) FACS-based cell ploidy assays for all cell lines. Singlets were isolated as in Figure 2D and used to plot a histogram of PI intensity for >9000 cells (number of cells, n, are above each histogram). Similar results were observed when the same experiment was performed on another independent occasion.

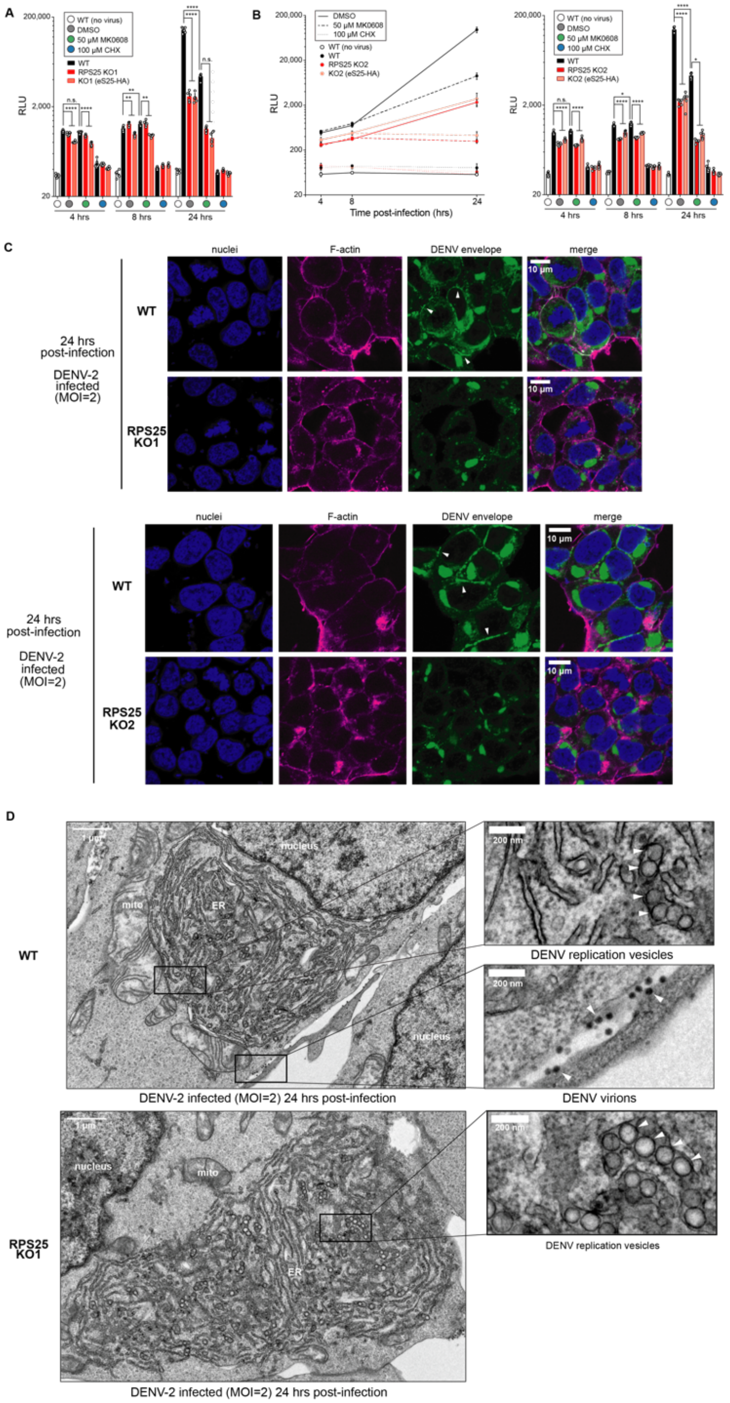


**Figure S8.** DENV resistance of the RPS25 KO acts at a late stage of infection when translation and replication are coupled. **A.** Full data for the time course of DENV-luc infection shown in Figure 4A, now showing individual data points and statistical significance for all time points. Error bars represent the 95% CI of n=6 biological replicates and data were analyzed with a two-way ANOVA with Tukey test. **B.** Time course of DENV-luc infection in WT, RPS25 KO2, and eS25-HA AB cells. Experiment performed for RPS25 KO2 cells as in Figure 4A. P-values: ≥0.05 (n.s.), 0.01-0.05 (*), 0.001-0.01 (**), and <0.0001 (****). **C.** Confocal images from IF staining of WT and RPS25 KO HAP1 cells infected with DENV-2 at MOI=2. The top two panel represent the full images set of images used in Figure 4C. White arrows point to bead-like structures of the E protein within the cell periphery of infected WT cells that is largely absent from KO cells. **D.** TEM images of WT and RPS25 KO HAP1 cell sections prepared following infections with DENV-2 at MOI=2. ER membrane-derived replication vesicles (top and bottom right) and virions are indicated with white arrows.

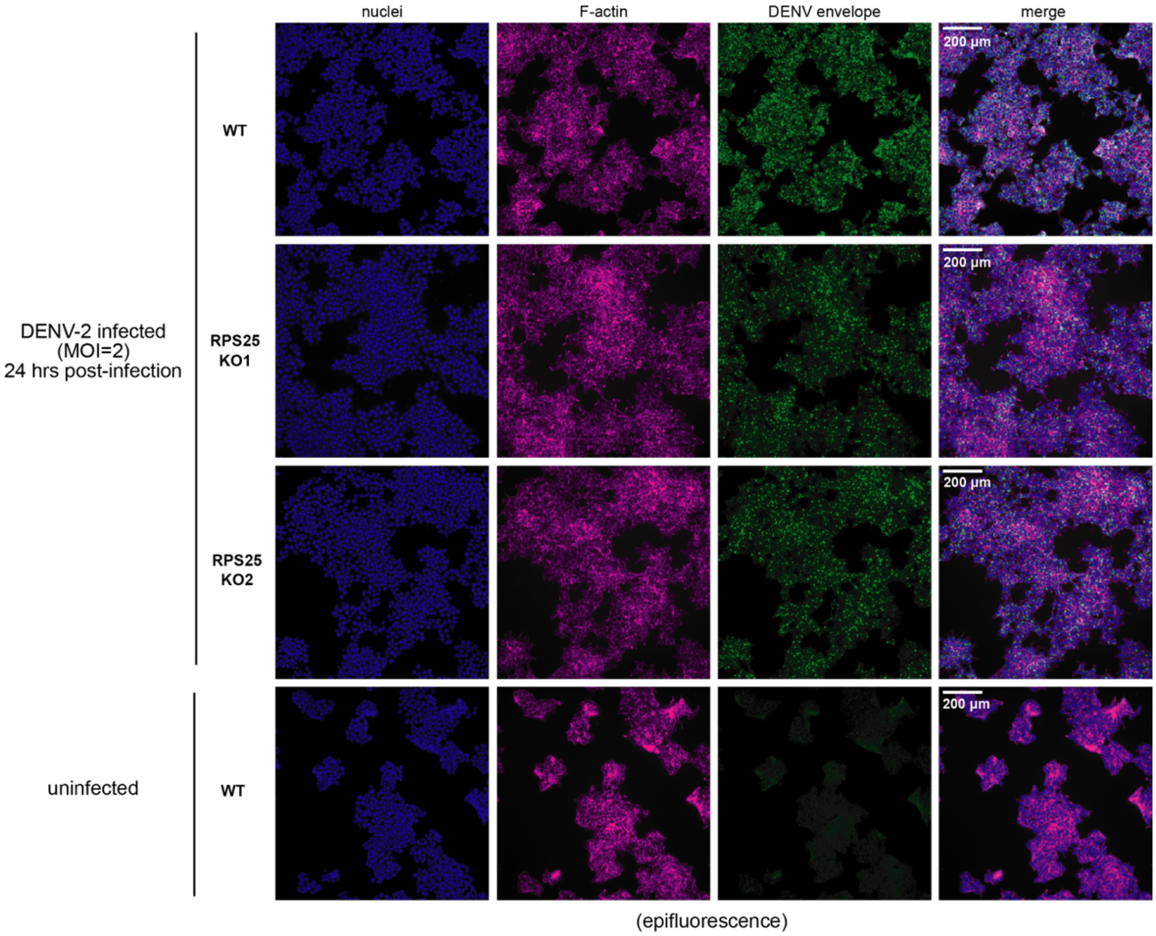


**Figure S9.** Epi-fluorescent imaging of fixed WT and RPS25 KO HAP1 cells for the DENV envelope protein following infection with DENV-2 at MOI=2. Same samples as in Figure 4C and Figure S8, showing widefield images so as to demonstrate the trend for staining across many cells. The uninfected control (bottom) is included to demonstrate the specificity of the antibody.

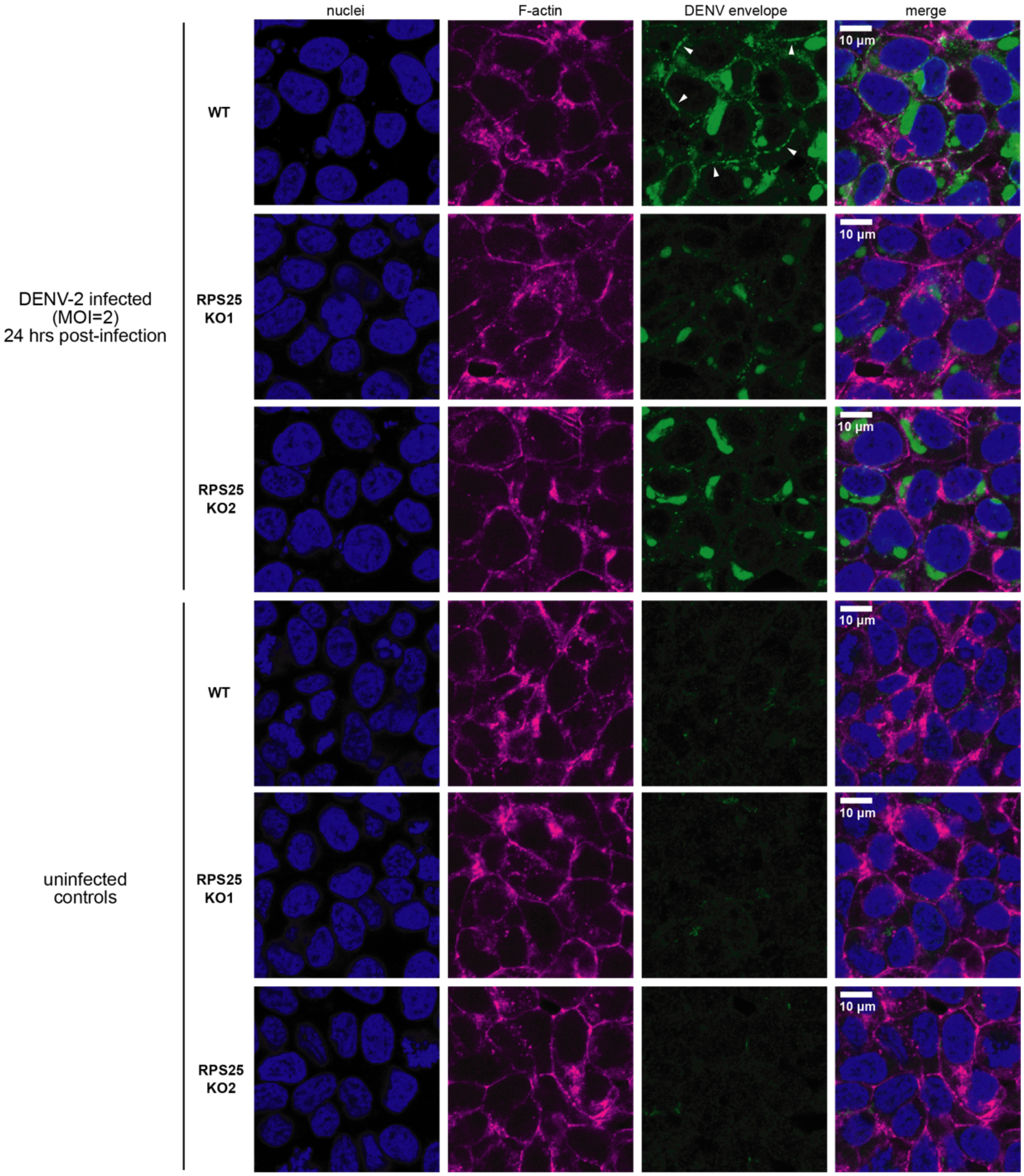


**Figure S10.** Additional confocal images of fixed HAP1 cells with IF staining for the envelope protein following DENV-2 infection, as in Figure 4C and Figure S8. Both infected (MOI=2) and uninfected cells are shown to demonstrate the specificity of the antibody, and white arrows point to bead like structures in WT cells.

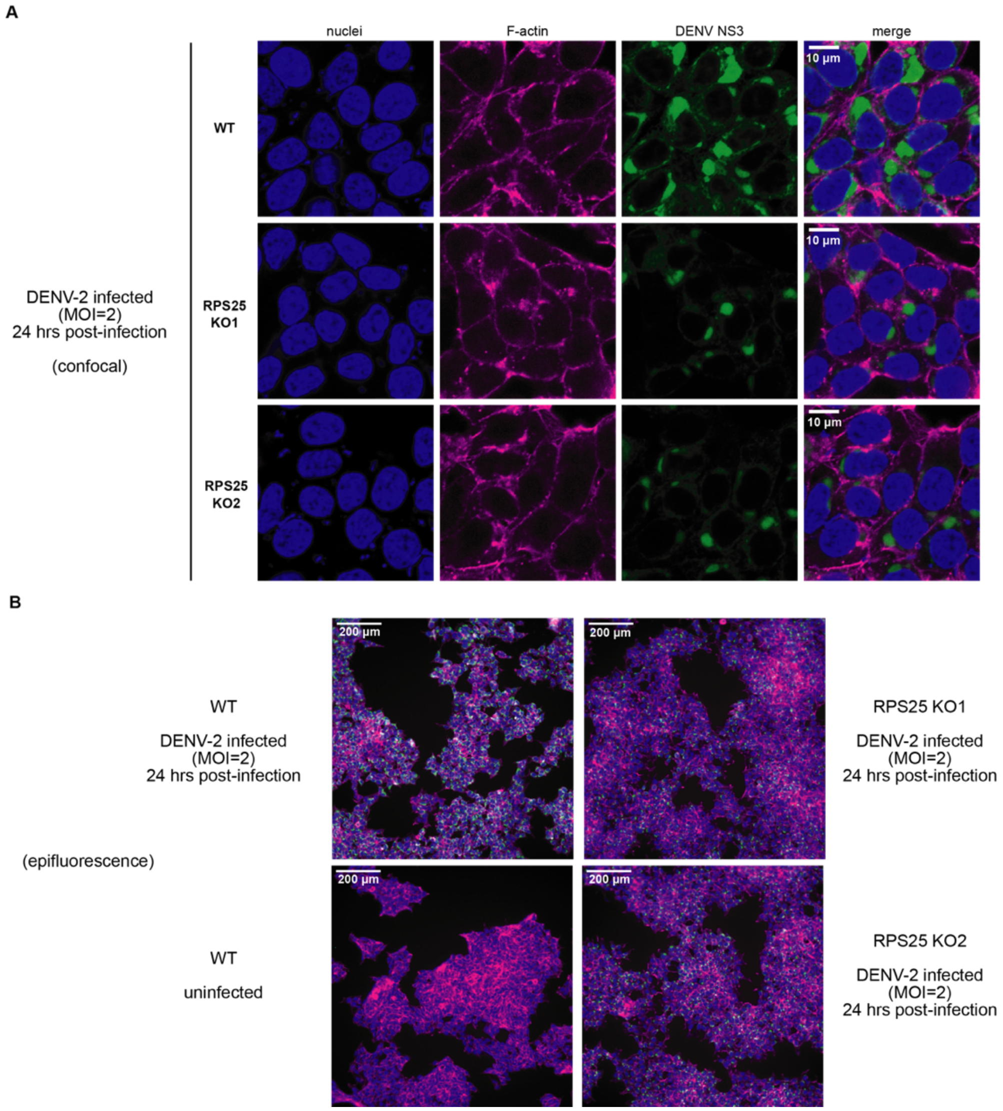


**Figure S11.** IF imaging of DENV-2 infected HAP1 cells (MOI=2) for the DENV NS3 protein. Fixed cells were stained with an antibody against the DENV NS3 protein alongside staining of nuclei with Hoescht and F-actin with Phalloidin 660. Unlike the staining for the DENV E protein, bead like structures are not observed at the cell periphery. **A.** Confocal images of DENV-2 infected WT and RPS25 KO HAP1 cells. **B.** Epifluorescent imaging of DENV-2 infected and uninfected HAP1 cells. Merged widefield images included alongside uninfected WT control to demonstrate specificity of NS3 staining. Nuclei, F-actin, and NS3 staining are colored the same as in (A).

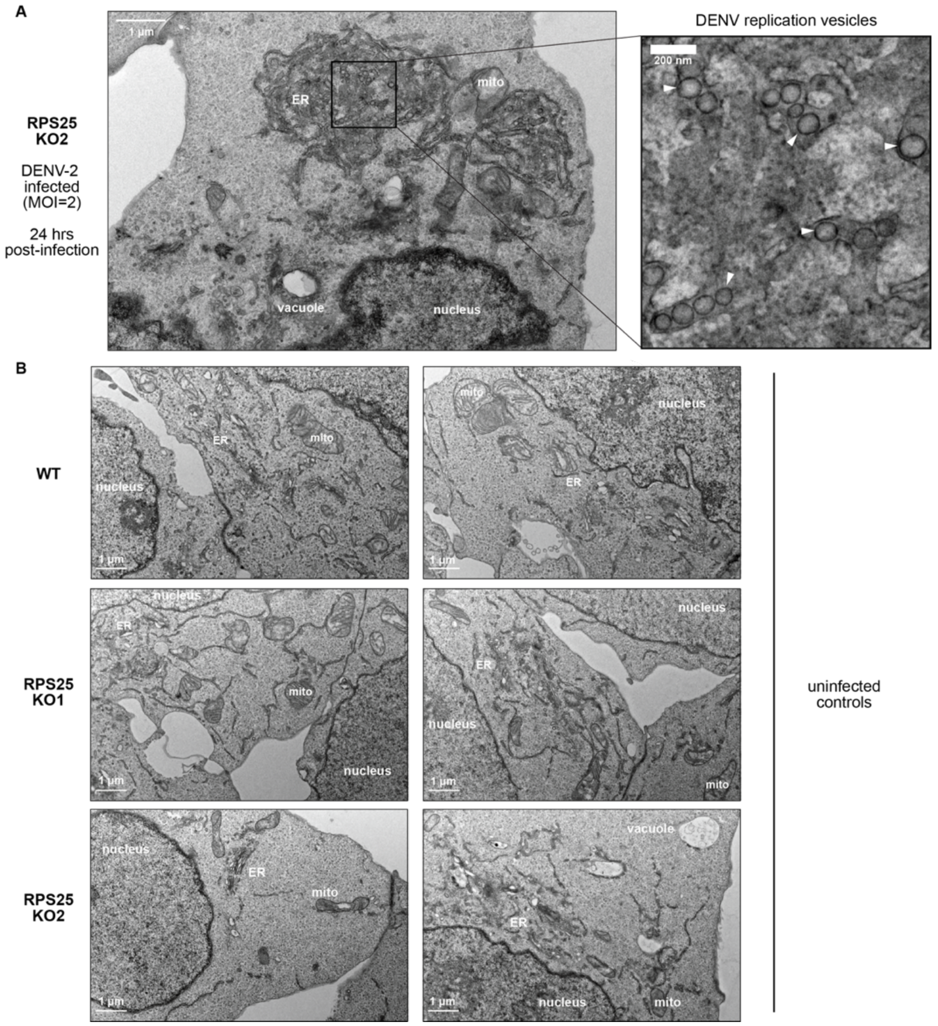


**Figure S12.** Transmission electron microscopy of WT and RPS25 KO HAP1 cells. **A.** A fixed and embedded cell slice from infection of RPS25 KO2 HAP1 cells with DENV-2 at MOI=2. As in Figure 4D, white arrows point to the appearance of ER membrane-derived replication vesicles within the inset on the right. **B.** Example cell slices from uninfected WT and RPS25 KO HAP1 cells, wherein no replication vesicle-like structures are observed.

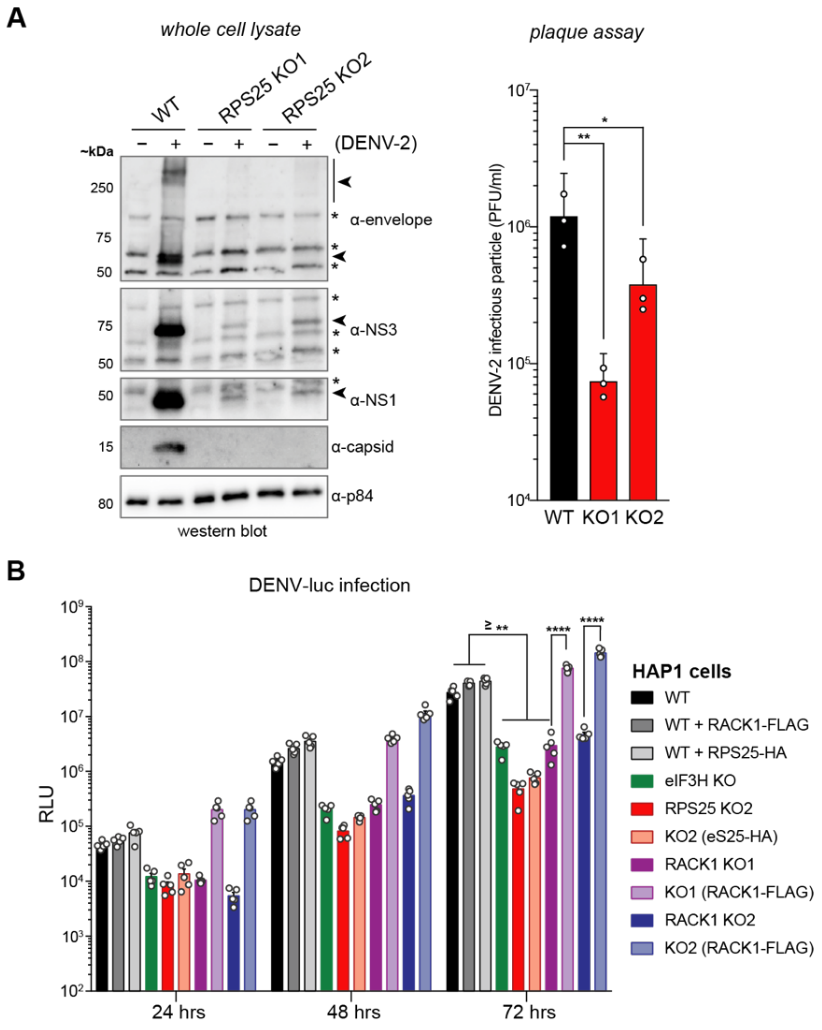


**Figure S13.** Viral infections of cells lines demonstrate that the RPS25 KO effect is robust and specific to the HAP1 cell lines and flaviviruses. **A.** Western blot analysis (left) and plaque assay (right) from infection with DENV-2 at MOI=0.1 and harvest at 48 hours post-infection. * indicates likely nonspecific protein bands. Plaque assay was performed using the virion-containing supernatant as described in the Methods section. The error bars in the plaque assay represent the 95% CI of three biological replicates. The number of infectious particles was compared between WT and KO cell lines using a one-way ANOVA with a Dunnett test to account for multiple comparisons. **B.** Additional example of a DENV-luc infection experiment assayed at 24, 48, and 72 hours post-infection with the DENV-luc virus at a MOI of 0.018 (similar to Figure 5B). Example shows that the RACK1 KO effect is consistent in two clonal KOs. Error bars represent the 95% CI of n=5 biological replicates and a one-way ANOVA was performed for each time point, correcting for multiple comparisons with a Tukey test. For simplicity, plot only shows significance values for 72 hrs post-infection. P-values: ≥ 0.05 (n.s.), 0.01-0.05 (*), 0.001-0.01 (**), 0.0001-0.001 (***), and <0.0001 (****).

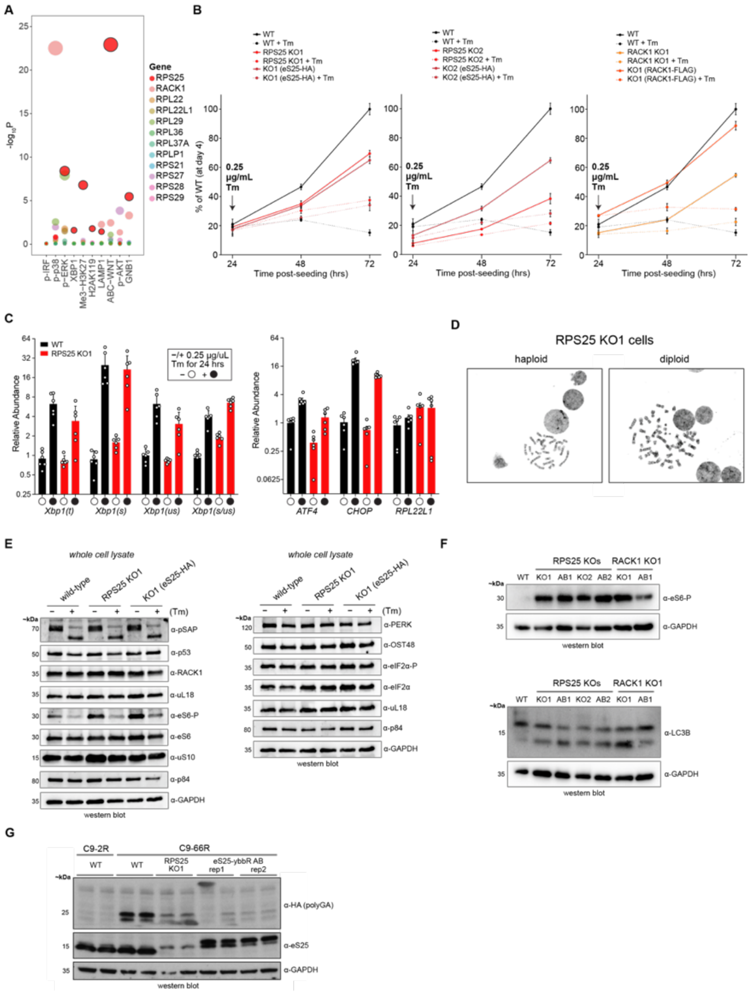


**Figure S14.** RPS25 loss drives pleiotropic phenotypes. **A.** Plot of significance scores (negative log10 FDR-corrected P-values, $-\log_{10}P$) for RP genes with low essentiality scores from ten published HAP1 FACS-based genetic screens (2, 3). **B.** MTT proliferation assays of WT HAP1 cells versus RPS25 KO1, RPS25 KO2, and RACK1 KO1 with their respective addbacks. Data from the left two panels represents the data that was modified for Figure 4E. Error bars for all panels represent the 95% CI of n=5 biological replicates. **C.** The RPS25 KO has a responsive UPR. RT-qPCR assays of WT and RPS25 KO1 assayed for Xbp1 splicing, other UPR markers (ATF4 and CHOP), and RPL22L1. Markers were normalized to 18S rRNA expression and y-axis expression is represented on a log2 scale. The left plot relative abundances are normalized to WT Xbp1(us) at 1, while the right panel relative abundance is normalized to WT ATF4 at 1. Error bars represent the 95% CI of n=5 biological replicates. **D.** Karyotyping of WT and RPS25 KO cells by the UCSF Cytogenetic Core Laboratory. Twenty single cells from each cell lines were evaluated. The upper microscope viewfields are representative images of haploid and diploid cells from the RPS25 KO1 cell line. Complete documentation is available upon request. **E.** Western blots of cellular lysates treated with or without 0.25 μg/mL Tm as in Figure 5G, now probed for other ribosomal proteins (RACK1, RPL5/uL18, RPS6/eS6, RPS6-P/eS6-P, RPS20/uS10), the p53 tumor suppressor, an STT3A-specific glycosylated protein (pSAP), an OST complex protein (OST48), another UPR maker (PERK), as well as phosphorylated and total alpha subunit of eIF2 (eIF2α-P and eIF2α). The blotting results indicate little evidence of ribosome level alterations in the RPS25 KO, stable levels of p53, intact OST and STT3A-specific glycosylation, and relatively unaltered PERK branch of the UPR. Part of this blot is in Figure 5E. **F.** Western blots of WT, KO, and AB HAP1 cell lines for eS6 phosphorylation and LC3B lipidation. **G.** Western blots of WT, RPS25 KO1, and KO1 (eS25-ybbR AB) transfected with RAN translation constructs encoding different numbers of C9orf72 (GGGGCC) repeats (C9-2R and C9-66R: 2 and 66 repeats, respectively). Technical duplicates were run in side-by-side lanes and two transfection replicates (rep1 and rep2) of the eS25-ybbR AB are shown. In these blots, the eS25 antibody detects a likely non-specific band in all samples that migrates above the expected eS25 molecular weight (MW). The eS25-ybbR AB construct was used instead of the eS25-HA AB construct due to interference of the eS25-HA tag with that of the HA-tagged polyGA.

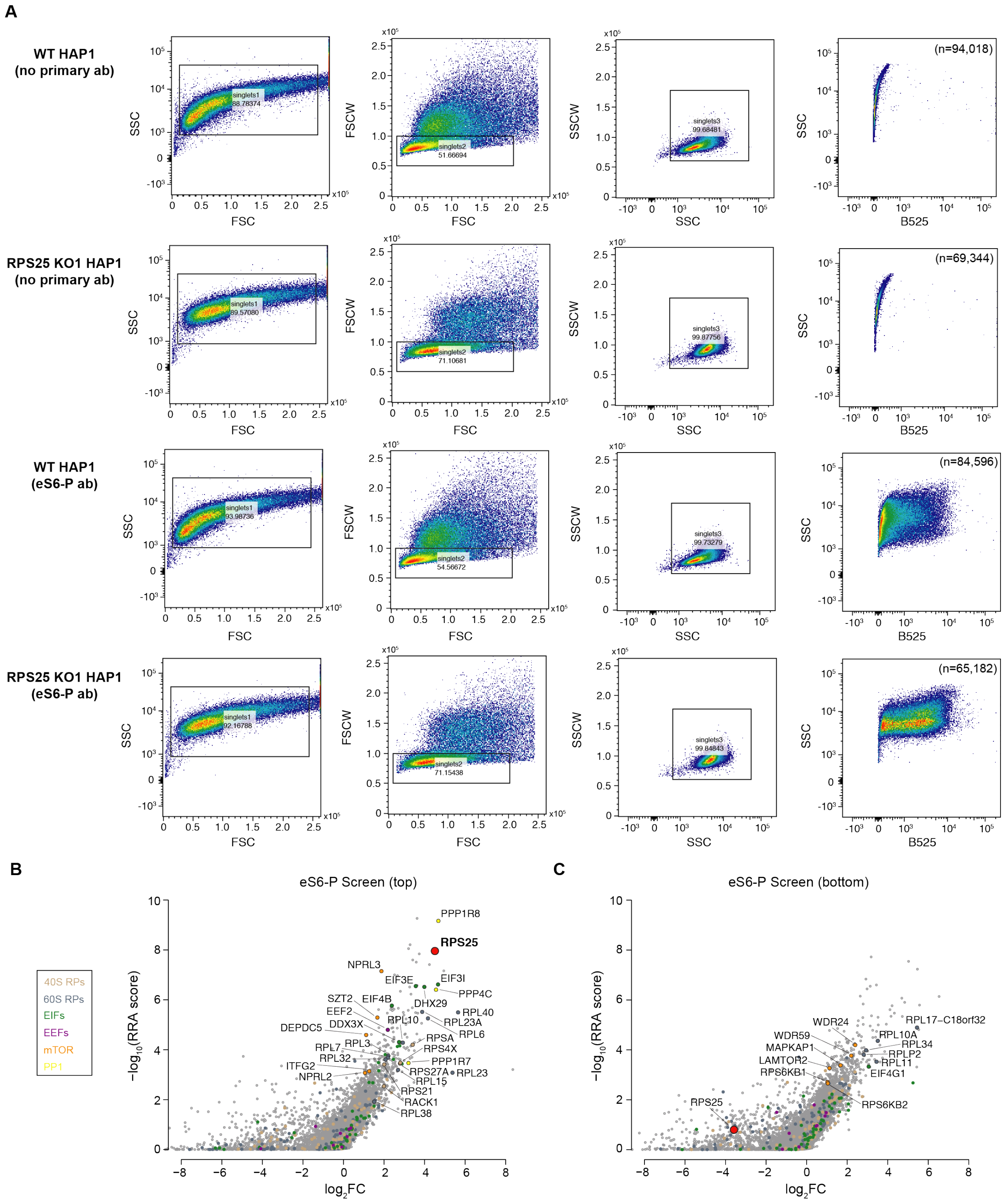


**Figure S15.** Genome-wide CRISPR-Cas9 screen for phosphorylated eS6 regulators. **A.** Immunoflow optimization of WT and RPS25 KO1 cells stained for eS6-P. Plots show the identification of singlets and the populations of antibody-stained cells (B525, Alexa 488 secondary antibody). No antibody controls included for completeness. **B.** Plots for score of screens in the positive selection (-$\log_{10}$(RRA score)) versus fold-change enrichment ($\log_2$FC) for the top and bottom populations of eS6-P stained cells (~2%). Genes are colored and annotated according to the legend.

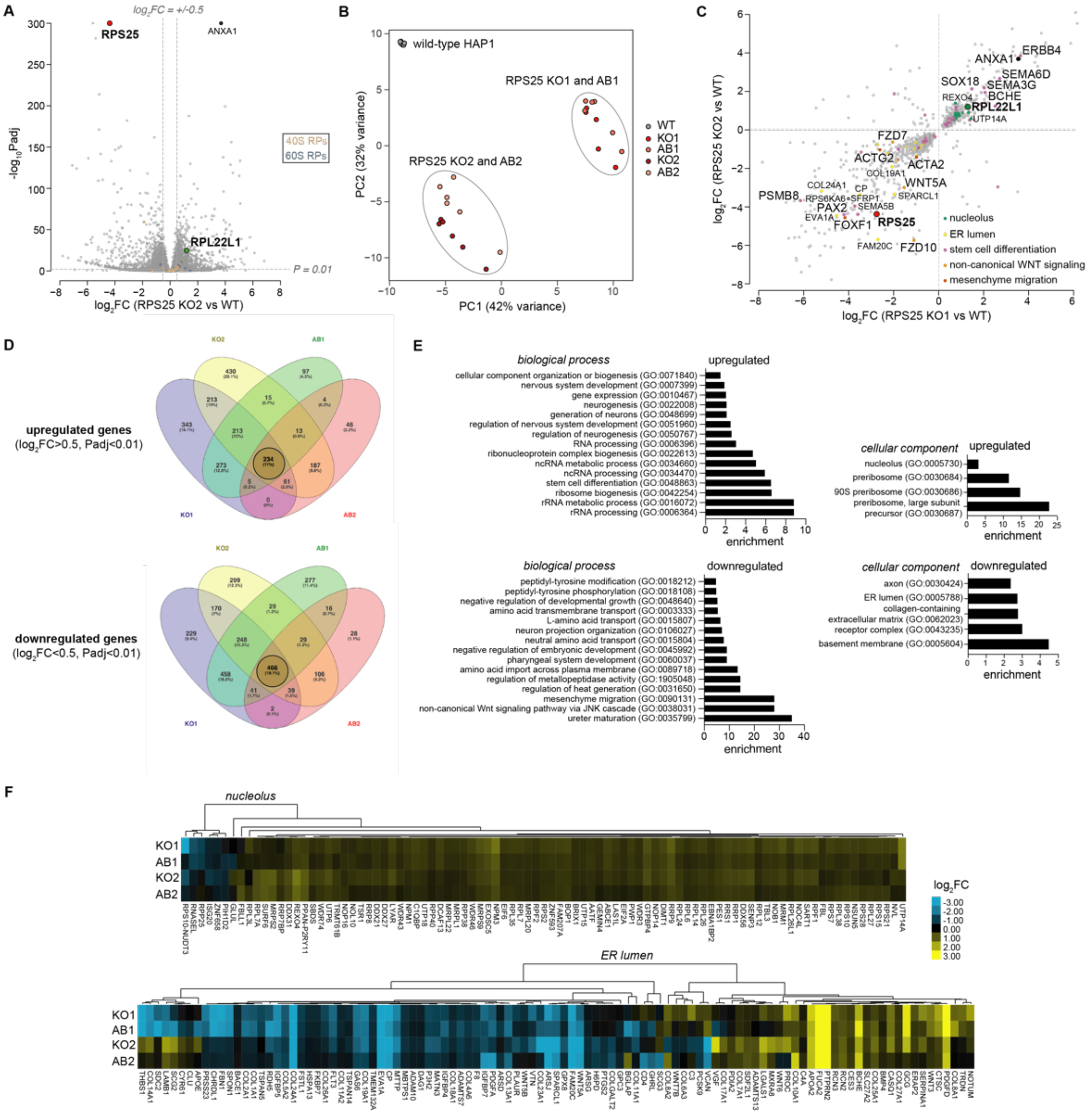

**Figure S16.** RNA-seq of WT, RPS25 KO, and eS25-HA AB cells demonstrates a shared transcriptional response from RPS25 loss. **A.** Volcano plot of significant fold-changes between RPS25 KO2 and WT HAP1 cells as in Figure 6A. **B.** Principal component analysis (PCA) of all samples from DESEQ2. Each bubble represents a biological replicate with n=6 per cell line. The percent variance for each principal component is indicated in the axis label. **C.** Common shared significant fold-changes for all RPS25 KOs and ABs. Showing FC for all genes with Padj<0.01 in RPS25 KO1 and KO2 conditions. Select genes are annotated based on categorization as indicated to highlight cellular compartments that match the ontology analysis, wherein upregulated nucleolus-related or downregulated ER lumen-related genes are colored. Select differentiation-related genes are also indicated, including those from stem cell differentiation, non-canonical WNT signaling, and mesenchyme migration ontologies. **D.** Venn diagrams for the shared fold-changes ($log_2FC$<-0.5 or >0.5, Padj<0.01) of both the RPS25 KOs and ABs ($log_2FC$<-0.5 or >0.5, Padj<0.01). Plot is similar to Figure 6B, but without proportional scaling. Shared genes for all conditions were used for ontology analysis. **E.** GO analysis showing fold enrichment for upregulated and downregulated genes. The top 15 GOs are shown (all with FDR-corrected P-value<0.05). **F.** Heatmaps from hierarchical clustering of $log_2FCs$ for each condition by genes within the nucleolus gene ontology (GO:0042254) and ER lumen gene ontology (GO:0005788). Only genes with $log_2FC$>0.5 in at least two conditions are shown.

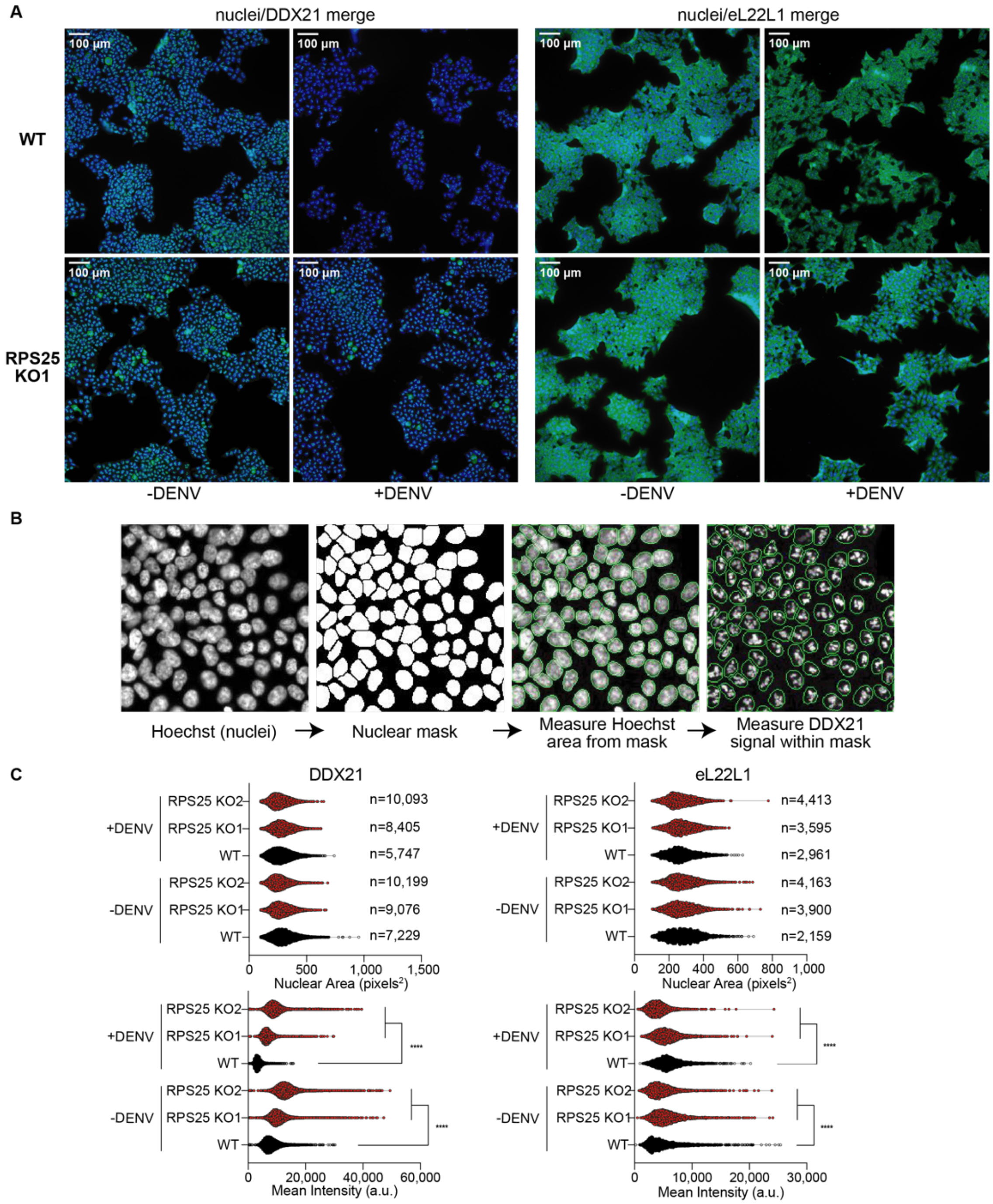


**Figure S17.** Quantitative image analysis of WT and RPS25 KO for nucleolar markers. **A.** Example IF images of WT and RPS25 KO cell lines stained for the nucleolar proteins DDX21 and eL22L1. Images are as in Figure S6. Only the images from epi-fluorescent imaging were used for analysis. Representative images are from WT and RPS25 KO1 HAP1 cell lines with or without (+/-) DENV at MOI=2, fixed and stained 24 hours post-infection. **B.** Example of automated nuclear boundary identification for use in image analysis in (D). Images are from viewfields of WT cells stained for Hoescht and DDX21 as in (A). **C.** Violin plots depicting parameters from image analysis of cell nuclei with antibody markers. DDX21 (left) and eL22L1 (right) mean intensity values (bottom panels) were determined by measurements from within nuclear regions (upper panels). For each cell line, the number of cells analyzed (n) is indicated next to bars in top upper panels. Part of this figure is replicated in Figure 6D. Statistical significance represents the results of a two-way ANOVA for each cell line and condition (-/+ DENV), correcting for multiple comparisons with a Tukey test. P-value: <0.0001 (****).

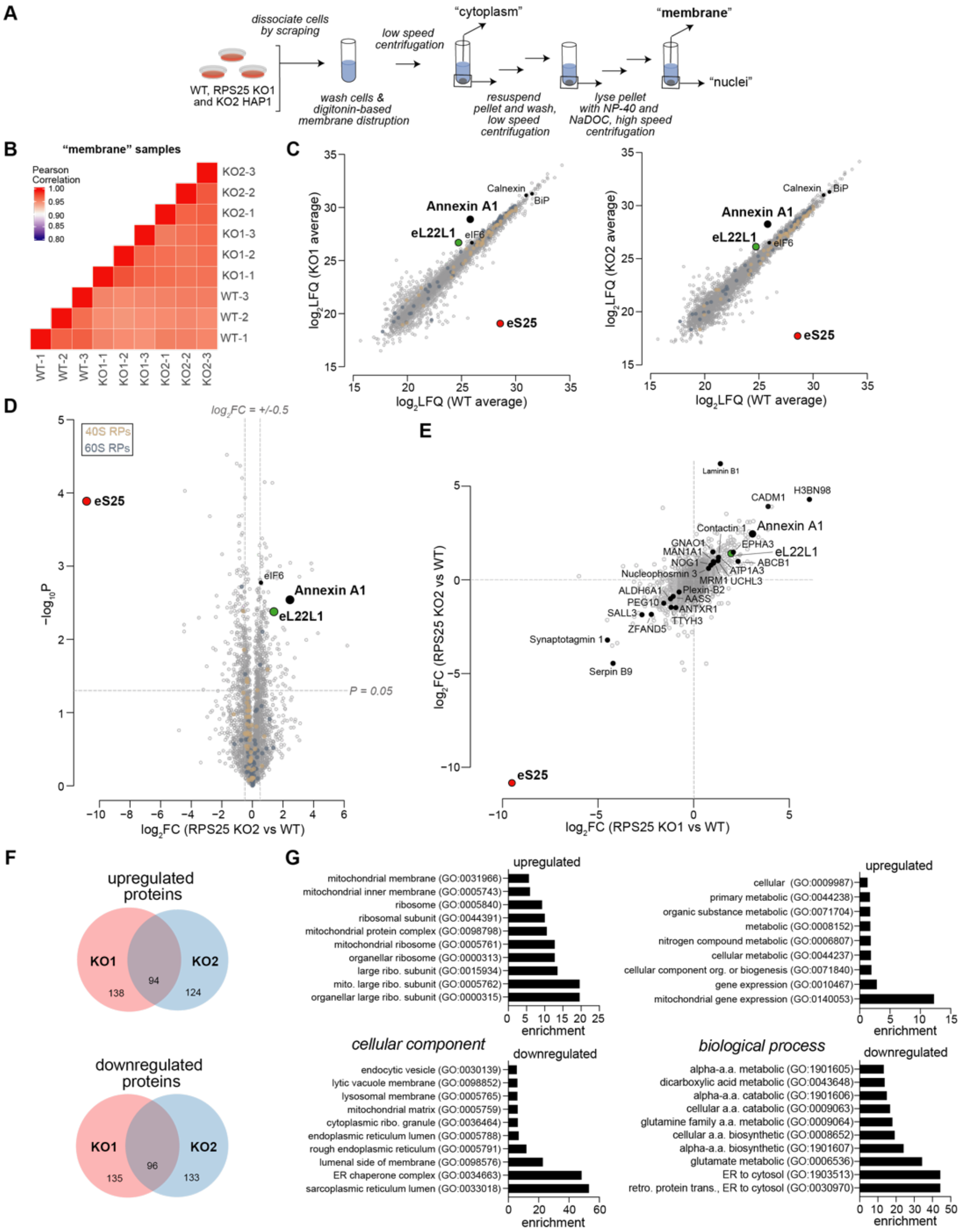


**Figure S18.** Membrane mass spectrometry of WT and RPS25 KO HAP1 cells. **A.** Schematic for purification of the "membrane" fraction from WT and RPS25 KO HAP1 cell lines. The membrane fraction as indicated was subsequently used for mass spectrometry analysis. **B.** Heatmap of Pearson correlation coefficients (ρ) for $log_2$LFQ intensities from each membrane sample. **C.** Average $log_2$LFQ values for RPS25 KO1 and KO2 versus WT membranes. Plot demonstrates the enrichment of ER proteins such as Calnexin and BiP in addition to other notable alterations. **D.** Volcano plot for significant fold-changes from LFQ intensities of membrane fractions. Plot as in Figure 7A. **E.** Common shared significant fold-changes for RPS25 KO1 and KO2 versus WT. **F.** Venn diagrams of upregulated and downregulated membrane proteins for RPS25 KO1 and KO2 versus WT samples. Proteins with $log_2FC < -0.5$ and $>0.5$, and $P<0.05$ were compared for KO1 and KO2 versus WT. **G.** Ontology analysis of changing membranes proteins with $log_2FC < -0.5$ and $>0.5$, and $P<0.05$. Analysis from cellular component (left) and biological process (right) ontology categories is shown. Fold-enrichment is shown for each ontology, selecting the top 10 (or less) with FDR-corrected P-values<0.05.

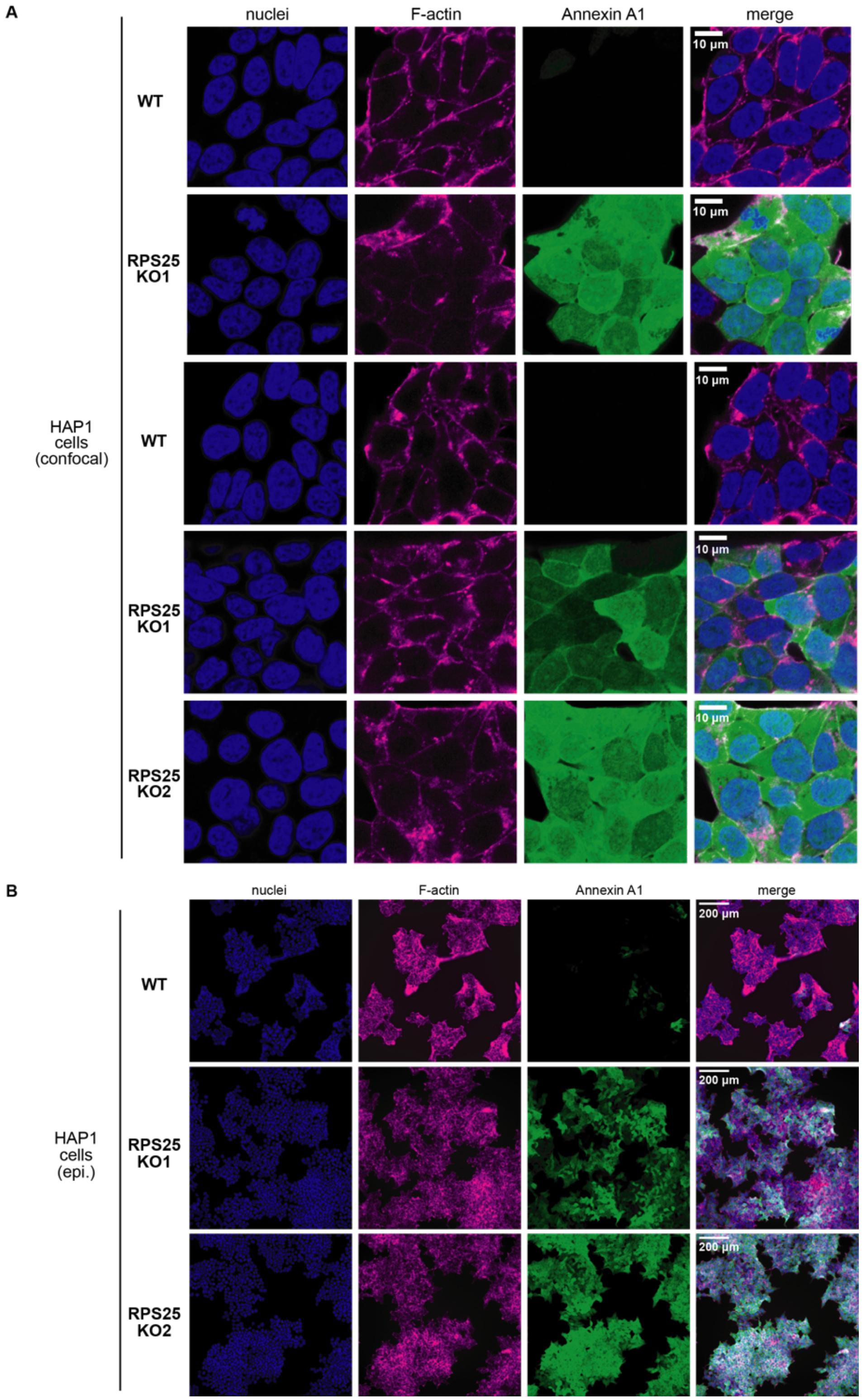


**Figure S19.** Immunofluorescent imaging of WT and RPS25 KO cells for Annexin A1. **A.** Confocal images of fixed WT and RPS25 KO HAP1 cells for Annexin A1 alongside staining of nuclei with Hoescht and F-actin with Phalloidin 660. Top two panels represent the complete images from Figure 7D. **B.** Epi-fluorescent imaging of WT and RPS25 KO HAP1 cells to demonstrate widefield images of the same samples imaged by confocal.

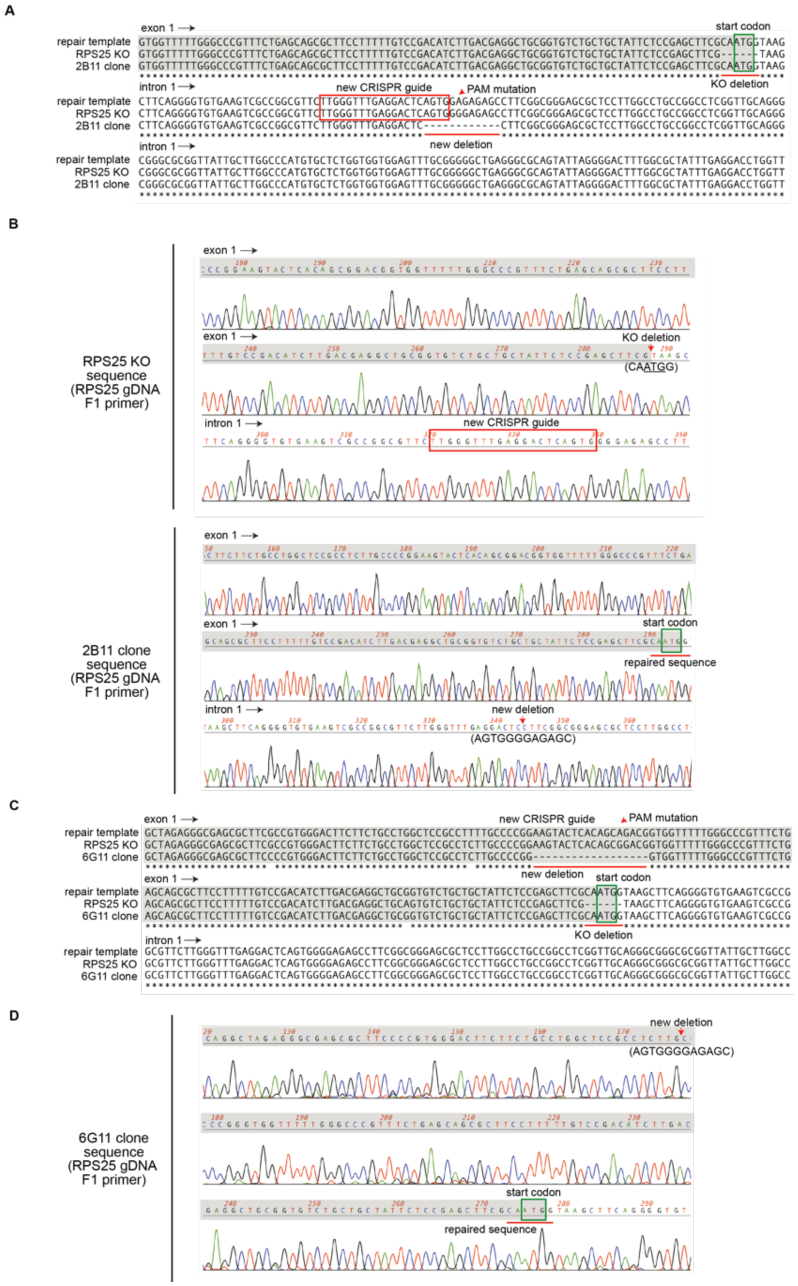


**Figure S20.** Homology-directed repair of the RPS25 KO1 genomic locus and sequence validation of the first (2B11) and second repair clone HDR2 (6G11). **A.** Sequence alignment of a region of the designed repair template with that of the of the RPS25 KO1 genomic sequence and the sequence of the HDR1 (2B11) clone. Alignment is annotated to show the exon (gray highlight) and intron (no highlight) as well as the start codon (green box) at the site of the original CRISPR/Cas9 deletion. **B.** Sequence chromatograms from sequencing the RPS25 KO1 cell line (top) and the RPS25 HDR repair cell line 2B11 (bottom). Sequences are from priming an amplified genomic DNA fragment with the RPS25 gDNA F1 primer, which initiates upstream at the 5' end of the RPS25 genomic locus. **C.** Sequence alignment of a region of the designed repair template with that of the of the RPS25 KO1 genomic sequence and the sequence of the HDR2 (6G11) clone. Alignment is annotated to show the exon (gray highlight) and intron (no highlight) as well as the start codon (green box) at the site of the original CRISPR/Cas9 deletion. **D.** Sequence chromatograms from sequencing the RPS25 KO1 cell line (top) and the RPS25 HDR repair cell line 6G11 (bottom). Sequences are from priming an amplified genomic DNA fragment with the RPS25 gDNA F1 primer, which initiates upstream at the 5' end of the RPS25 genomic locus. Regions of interest are annotated for chromatograms to indicate modified sites in the context of the exon and intron sequences.

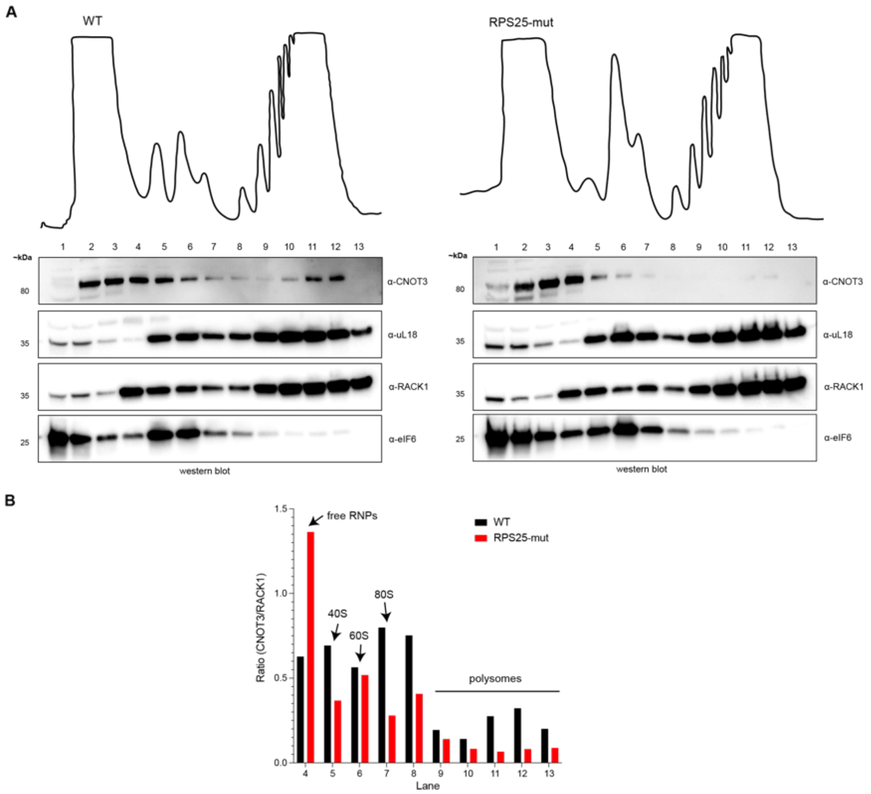


**Figure S21.** Polysome profiles of wild type (WT) and N-terminal RPS25 mutant (RPS25-mut) HEK293T cells. **A.** Polysome profile traces from WT and RPS25-mut HEK293T cells. Post-nuclear lysate was sedimented in 10-60% sucrose gradients, the top and bottom of the gradients are indicated, and the height of peaks is proportional to the absorbance at 254 nm. Fractions were probed by immunoblotting for select ribosomal proteins (RACK1, uL18), the 60S-binding protein eIF6, and the CCR4-Not subunit CNOT3. Human CNOT3 is 22% identical to the *S. cerevisiae* Not5 subunit that interacts with eS25's N-terminus. **B.** Relative densitometry of CNOT3 (normalized to RACK1) in polysome profile fractions.
